# Structural basis for the allosteric regulation of the SbtA bicarbonate transporter by the P_II_-like protein, SbtB, from *Cyanobium* sp. PCC7001

**DOI:** 10.1101/762807

**Authors:** Joe A. Kaczmarski, Nan-Sook Hong, Bratati Mukherjee, Laura T. Wey, Loraine Rourke, Britta Förster, Thomas S. Peat, G. Dean Price, Colin J. Jackson

## Abstract

Cyanobacteria have evolved a suite of enzymes and inorganic carbon (C_i_) transporters that improve photosynthetic performance by increasing the localized concentration of CO_2_ around the primary CO_2_-fixating enzyme, Rubisco. This CO_2_-concentrating mechanism (CCM) is highly regulated, responds to illumination/darkness cycles and allows cyanobacteria to thrive under limiting C_i_ conditions. While the transcriptional control of CCM activity is well understood, less is known about how regulatory proteins might allosterically regulate C_i_ transporters in response to changing conditions. Cyanobacterial sodium-dependent bicarbonate transporters (SbtAs) are inhibited by P_II_-like regulatory proteins (SbtBs), with the inhibitory effect being modulated by adenylnucleotides. Here, we used isothermal titration calorimetry to show that SbtB from *Cyanobium* sp. PCC7001 (SbtB7001) binds AMP, ADP, cAMP and ATP with micromolar-range affinities. X-ray crystal structures of apo- and nucleotide-bound SbtB7001 revealed that while AMP, ADP and cAMP have little effect on the SbtB7001 structure, binding of ATP stabilizes the otherwise flexible T-loop and that the flexible C-terminal C-loop adopts several distinct conformations. We also show that ATP binding affinity is increased ten-fold in the presence of Ca^2+^ and we present an X-ray crystal structure of Ca^2+^ATP:SbtB7001 that shows how this metal ion facilitates additional stabilizing interactions with the apex of the T-loop. We propose that the Ca^2+^ATP-induced conformational change observed in SbtB7001 is important for allosteric regulation of SbtA activity by SbtB and is consistent with changing adenylnucleotide levels in illumination/darkness cycles.

**GRAPHICAL ABSTRACT:** 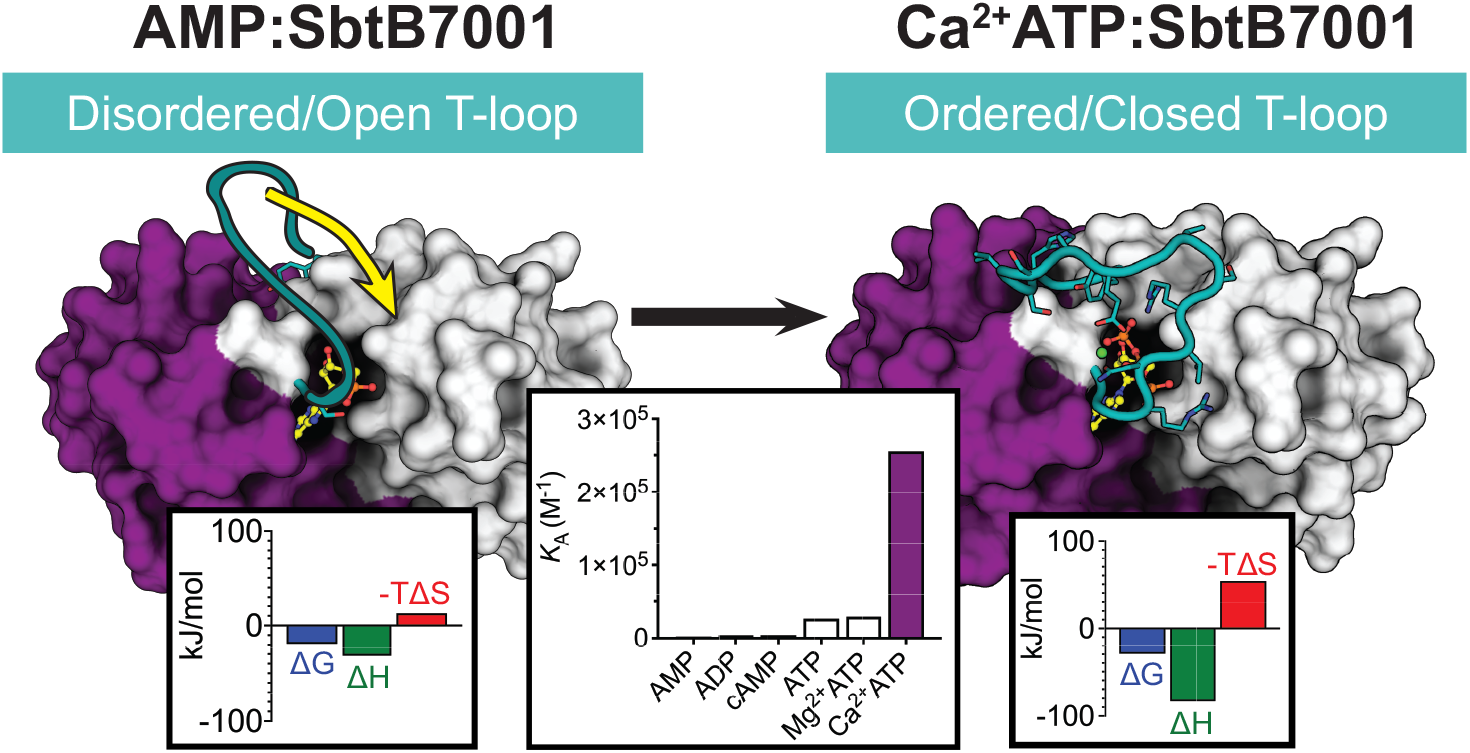

## 1. Introduction

Cyanobacteria possess a highly regulated network of enzymes, inorganic carbon (C_i_) transporters and specialized compartments that enhance photosynthetic performance by increasing the localized concentration of CO_2_ around the primary CO_2_-fixating enzyme, Rubisco [1,2]. This CO_2_-concentrating mechanism (CCM) responds to illumination/darkness cycles and allows cyanobacteria to thrive under limiting C_i_ conditions [3]. Bicarbonate (HCO_3_^−^) transporters, which maintain high concentrations of HCO_3_^−^ in the cell (up to 40 mM), are of vital importance to the CCM [3]. While the transcriptional regulation of these HCO_3_^−^ transporters has been characterized [4], the molecular signals and post-translational mechanisms that transiently control the activity of cyanobacterial HCO_3_^−^ transporters remain unclear.

The high-affinity, low-flux, sodium-dependent bicarbonate transporter A (SbtA), which is found in the plasma membrane of many cyanobacterial species, is a key component of the cyanobacterial CCM [3]. The dicistronic operon that typically encodes both SbtA and a cytoplasmic P_II_-like regulatory protein (SbtB) is upregulated under C_i_-limiting conditions in *Synechocystis* and *Synechococcus* species [5–8]. We have previously shown that SbtB from *Synechococcus elongatus* PCC7942 (SbtB7942) forms a stable complex with SbtA7942, and that several SbtB homologs from *Synechococcus, Synechocystis* and *Cyanobium* sp. inhibit their cognate SbtA when coexpressed in *E. coli* [9]. Since SbtA-mediated HCO_3_^−^ transport must be coupled with the concurrent activity of ATP-dependent sodium exporters, we proposed that SbtB limits futile cycling by directly regulating SbtA-mediated HCO_3_^−^ transport in cyanobacteria through the formation of an inhibitory SbtA-SbtB complex [9]. This general model was recently confirmed by Selim *et al.* [10], who showed that SbtB from *Synechocystis* sp. PCC6803 (SbtB6803) forms a stable complex with membrane-bound SbtA6803 under limiting C_i_ conditions *in vivo* and responds to changes in the relative levels of adenylnucleotides in the cell.

SbtB is a non-canonical member of the P_II_ signal-transduction superfamily [9–11]. P_II_ proteins are homotrimeric proteins that typically sense and tune the metabolic state of cells by binding adenylnucleotides and small molecules and forming regulatory complexes with diverse protein targets including transcription factors, enzymes and membrane transporters [12]. The functions of P_II_ proteins are primarily driven by large, ligand-induced conformational changes in the three flexible, solvent-exposed T-loops that mediate interactions with target proteins. Competitive binding of adenylnucleotides (typically ADP/Mg^2+^ATP) and/or other effector molecules at the clefts between the subunits can alter the conformation of the T-loops and modulate the affinity of the P_II_ protein for its target [13].

Selim *et al.* [10] recently reported that SbtB6803 binds AMP, ADP, cyclic-AMP (cAMP) and ATP with micromolar-range affinities, although the reported range of *K*_D_ values was relatively narrow (11–46 μM from microscale thermophoresis experiments, 75–250 μM for *K*_D3_ from ITC experiments). AMP and ADP were observed to stabilize the SbtA–SbtB6803 complex *in vivo*, and the authors suggested that the cAMP:AMP ratio may act as a signal to control SbtA6803-mediated HCO_3_^−^ transport *in vivo*. However, the structures of SbtB6803 in complex with AMP and cAMP did not show any ligand-induced T-loop rearrangements that are typically required for the differential binding of P_II_ proteins to their target proteins [10]. Therefore, the structural basis for SbtA-inhibition by the formation of the SbtA–SbtB complex remains unclear.

Furthermore, the functional consequence of sequence differences among SbtB homologs have not been explored. For example, many SbtB homologs, such as SbtB from *Cyanobium* sp. PCC7001 (SbtB7001), lack the C-terminal extension that forms a putative redox-sensing domain in SbtB6803 [10]. Additional structural and functional data on phylogenetically distinct SbtB homologs are required to achieve a deeper understanding of the molecular basis for the regulation of CCM activity, *via* regulation of SbtA-mediated HCO_3_^−^ transport, by SbtB.

Here we show that although SbtB7001 binds AMP, ADP, ATP and cAMP with micromolar-range affinities (like SbtB6803 [10]), the presence of Ca^2+^ dramatically increases the affinity of SbtB7001 for ATP affinity such that is ~50-fold and 100-fold greater than the affinity for ADP/cAMP and AMP (in the absence of Ca^2+^), respectively. High resolution crystal structures reveal that AMP, ADP and cAMP have little effect on the structure of SbtB7001, while ATP/Ca^2+^ATP binding causes dramatic rearrangements in the structure of SbtB7001, providing a molecular mechanism for the allosteric modulation of SbtB7001 function by ligand binding.

## 2. Materials and Methods

### 2.1 Cloning, expression and purification of SbtB7001

The gene encoding SbtB7001 (CPCC7001_1671) was codon-optimized for expression in *E. coli*, synthesized and cloned into pUC57 by GenScript, and then sub-cloned into the pHUE vector [14] between BamHI and HindIII sites to yield pHUE-SbtB7001. *E. coli* strain DH5α cells were used for cloning. pHUE adds polyhistidine and ubiquitin tags (His_6_–Ub) to the N-terminus of SbtB7001. Successful cloning was confirmed by Sanger sequencing at Garvan Molecular Genetics.

Protein expression was carried out in BL21(DE3) cells. Briefly, cells were grown in autoinduction media (20 g/l tryptone, 5 g/l yeast extract, 5 g/l NaCl, 6 g/l Na_2_HPO_4_, 3 g/l KH_2_PO_4_, 6 ml/l glycerol, 2 g/l lactose, 0.5 g/l glucose, 100 mg/l ampicillin), and protein expression was carried out overnight at 25 °C with shaking. His_6_–Ub-tagged SbtB7001 was purified from cell lysate using a 5 ml HisTrap FF column (GE Healthcare): the pre-equilibrated and lysate-loaded column was washed with binding buffer (20 mM NaH_2_PO_4_, 500 mM NaCl, 20 mM imidazole, pH 8), non-specific binders were removed using a step of 7% elution buffer, and tagged-SbtB7001 was eluted using 100% elution buffer (20 mM NaH_2_PO_4_, 500 mM NaCl, 500 mM imidazole, pH 8.0). Fractions containing tagged-SbtB7001 were pooled and transferred into binding buffer using a HiPrep 26/10 Desalting column (GE Healthcare). The His_6_–Ub tags were removed using a ubiquitin carboxyl-terminal hydrolase 2 catalytic core domain (USP2cc):SbtB7001 molar ratio of 1:25 at 4 °C overnight as previously described [14]. Untagged SbtB7001 was recovered using a 5 ml HisTrap FF column (GE Healthcare) using binding buffer and concentrated using a 3 kDa high molecular weight cut-off Amicon Ultra Centrifugal Filter (Merck). SDS-PAGE was used to assess the purity of untagged SbtB7001. For preparative purification, untagged SbtB7001 was loaded onto a HiLoad Superdex 75 16/600 size exclusion chromatography (SEC) column (GE Healthcare), and eluted in SEC buffer (20 mM HEPES, 150 mM NaCl, pH 8.0). For analytical size-exclusion, untagged SbtB7001 was loaded on a Superdex 75 10/300 GL column (GE Healthcare) and eluted in SEC buffer. The size-exclusion column was calibrated using a set of standard proteins (Gel Filtration HMW Calibration Kit, GE Healthcare) in SEC buffer. Protein concentration was measured spectrophotometrically using the molar absorption coefficient for USP2-cleaved SbtB7001 (12950 M^−1^ cm^−1^) as calculated using ProtParam (http://expasy.org/tools/protparam.html). SEC-purified, untagged SbtB7001 was used for isothermal titration calorimetry and crystallization.

### 2.2 Isothermal Titration Calorimetry

All isothermal titration calorimetry (ITC) experiments were performed on a Nano-ITC low-volume calorimeter (TA Instruments). ITC experiments were performed at 25 °C with stirring at 350 rpm. Protein and ligand solutions were prepared in matched SEC buffer and degassed before use. Solutions of HCO_3_^−^, AMP, ADP, ATP, and cAMP (all sodium salts, Sigma) were prepared volumetrically. Titrations were performed using 100–400 μM (monomer concentration) of USP2-cleaved SbtB7001, and typically involved 1 × 1 μl, followed by 22 × 2 μl injections of 2– 25 mM ligand with 250–300 s intervals between injections. Data from each titration were analyzed using NITPIC [15] and SEDPHAT [16]; the baseline-subtracted power was integrated, and the integrated heats were fit to the single binding site model (A + B <–> AB, hetero-association) model to obtain the macroscopic association constant (*K*_a_) and enthalpy of binding (ΔH). Values for the incompetent fraction of protein were constrained between −0.2 and 0.2 and were fitted locally for each experiment, as suggested in the documentation for SEDPHAT [16]. Global fitting was achieved by iteratively cycling between Marquardt–Levenberg and Simplex algorithms in SEDPHAT until modelling parameters converged. Model parameters were obtained through the global fitting of single titrations or replicates. 68.3% confidence intervals were calculated using the automatic confidence interval search with projection method using F-statistics in SEDPHAT. Other binding parameters and associated confidence intervals were obtained by propagation.

### 2.3 Protein crystallization

High throughput, sitting-drop vapor diffusion crystallization screens and subsequent hanging-drop optimization screens were set up at 18 °C or 4 °C in-house or at the Collaborative Crystallisation Centre (C3, CSIRO) and crystals formed in a number of conditions. For apo-SbtB7001 crystals, SEC-purified, untagged SbtB7001 was transferred to MES crystallization buffer (50 mM MES, pH 6.0, 100 mM NaCl). Crystals in sitting drops at 18 °C containing 0.15 μl of 10 mg/ml protein solution and 0.15 μl of reservoir solution (0.025 M disodium hydrogen-potassium dihydrogen phosphate, 20 % PEG 3350) were cryo-protected with 30 % (v/v) 1:1 mix of glycerol:ethylene glycol. Crystals of AMP:SbtB7001 formed at 18 °C in hanging drops containing 1 μl of protein solution (22 mg/ml of untagged SbtB7001 in 20 mM HEPES, 100 mM NaCl, pH 8.0, 10 mM MgCl_2_, 12 mM AMP, 50 mM NaHCO_3_) and 1 μl of reservoir solution (0.2 M sodium acetate, 18% PEG 3350). These crystals were cryo-protected using 30% (v/v) 2-methyl-2,4-pentanediol (MPD) and 100 mM NaHCO_3_. Crystals of ADP:SbtB7001 formed at 18 °C in sitting drops containing 0.15 μl of protein solution (22 mg/ml of untagged SbtB7001 in 20 mM HEPES, 100 mM NaCl, pH 8.0, 10 mM MgCl_2_, 12 mM ADP, 50 mM NaHCO_3_) and 0.15 μl of reservoir solution (2.5 M ammonium sulfate). Crystals were cryoprotected with 30% (v/v) MPD and 100 mM NaHCO_3_. Crystals of cAMP:SbtB7001 formed at 18 °C in hanging drops containing 1 μl of protein solution (22 mg/ml of untagged SbtB7001 in 20 mM HEPES, 100 mM NaCl, pH 8.0, 10 mM MgCl_2_, 12 mM cAMP, 50 mM NaHCO_3_) and 1 μl of reservoir solution (0.2 M potassium acetate, 16% PEG 3350). Crystals were cryo-protected with 30% (v/v) MPD and 100 mM NaHCO_3_. Crystals of ATP:SbtB7001 formed at 18 °C in sitting drops containing 0.15 μl of protein solution (22 mg/ml of untagged SbtB7001 in 20 mM Tris, 100 mM NaCl, pH 8.0, 10 mM MgCl_2_, 12 mM ATP, 20 mM sodium bicarbonate) and 0.15 μl of reservoir solution (0.2 M lithium chloride, 20 % PEG 6000, 100 mM sodium acetate-acetic acid, pH 5). Crystals were cryo-protected with 30 % (v/v) glycerol. Crystals of Ca^2+^ATP:SbtB7001 formed at 18 °C in sitting drops containing 0.15 μl protein solution (20 mg/ml of untagged SbtB7001 in 20 mM Tris, 100 mM NaCl, pH 8.0, 10 mM MgCl_2_, 12 mM ATP, 20 mM NaHCO_3_) and 0.15 μl of reservoir solution (30 % MPD, 0.02 M CaCl_2_, 0.1 M sodium acetate-acetic acid pH 4.6). Crystals were cryoprotected with 30 % (v/v) MPD.

All crystals were flash-cooled in liquid nitrogen. X-ray diffraction data were collected at the MX2 beamline at The Australian Synchrotron. Data was processed using XDS [17] and Aimless [18], and molecular replacement was performed using either MolRep [19] or Phaser [20]. Chain A of apo-SbtB6803 (PDB: 5O3P, 1.7 Å) was used for molecular replacement of Ca^2+^ATP:SbtB7001. All other structures were solved by molecular replacement using the resulting Ca^2+^ATP:SbtB7001 structure. Iterative cycles of manual model building and refinement were performed using Coot 0.8.2 [21], refmac [22], and/or *phenix.refine* [23]. TLS parameter refinement was also used, using TLS groups automatically selected by *phenix.refine*. Anisotropic B-factor refinement (all atoms except water) was used for refinement of Ca^2+^ATP:SbtB7001 (1.04 Å) and ADP:SbtB7001 (1.42 Å). Models were optimized using the PDB_REDO server [24]. Data collection and refinement statistics are provided in Table 2.

## 3. RESULTS

### 3.1 Nucleotide binding to SbtB7001

We used isothermal titration calorimetry (ITC) to investigate the affinity of SbtB7001 for the proposed P_II_ protein ligands AMP, ADP, cAMP and ATP (Table 1, Figure 1). Titrations with AMP (Figure 1A) gave a macroscopic dissociation constant (*K*_D_, averaged across the three binding sites in the homotrimer) of approximately 450 μM, with an enthalpic contribution to binding of ΔH = −32 kJ/mol and an entropic cost of TΔS = −12.7 kJ/mol. Titrations with ADP (Figure 1B) and cAMP (Figure 1C) gave intermediate *K*_D_ values of 224 μM and 238 μM, respectively, with similar enthalpic (ΔH of −44 kJ/mol for ADP, −61 kJ/mol for cAMP) and entropic contributions (TΔS of −23.3 kJ/mol for ADP, of −40.1 kJ/mol for cAMP) to binding. In contrast, ATP showed the highest binding affinity with a *K*_D_ of approximately 37 μM (Figure 1D). The binding of ATP was associated with a much larger change in enthalpy (ΔH of −81.1 kJ/mol), indicative of the formation of numerous additional electrostatic interactions between ATP and SbtB7001. In addition, ATP binding resulted in a substantial loss in entropy (TΔS of −55.9 kJ/mol), which is consistent with the stabilization of a mobile region of the protein upon binding of ATP.

**Table 1.** Parameters for the binding of effector molecules to SbtB7001, measured by isothermal titration calorimetry (ITC).*

| | $K_D$ (μM) | $\Delta H$ (kJ/mol) | $T\Delta S$ (kJ/mol) |
| --- | --- | --- | --- |
| **AMP** (n=3) | **447**<br>(398, 502) | **-31.8**<br>(-36.2, -29.5) | **-12.7**<br>(-17.4, -10.14) |
| + 500 μM $\text{CaCl}_2$ (n=2) | **127**<br>(107, 150) | **-30.6**<br>(-41.2, -24.8) | **-8.40**<br>(-19.4, -2.24) |
| **ADP** (n=4) | **244**<br>(213, 264) | **-43.9**<br>(-46.1, -35.1) | **-23.3**<br>(-25.7, -14.2) |
| + 500 μM $\text{CaCl}_2$ (n=2) | **270**<br>(221, 301) | **-49.3**<br>(-54.1, -37.7) | **-29.0**<br>(-34.0, -16.8) |
| **cAMP** (n=4) | **238**<br>(213, 266) | **-60.8**<br>(-65.6, -57.6) | **-40.1**<br>(-45.2, -36.6) |
| + 500 μM $\text{CaCl}_2$ (n=2) | **144**<br>(140, 147) | **-67.6**<br>(-68.1, -65.8) | **-45.7**<br>(-46.3, -43.8) |
| **ATP** (n=5) | **37.7**<br>(33.2, 43.5) | **-81.1**<br>(-86.3, -76.3) | **-55.9**<br>(-61.4, -50.8) |
| + 10 mM $\text{MgCl}_2$ (n=2) | **34.3**<br>(19.2, 56.5) | **-79.0**<br>(-89.6, -70.2) | **-53.5**<br>(-65.4, -43.3) |
| + EDTA (n=1) | **52.0**<br>(44.2, 61.0) | **-90.3**<br>(-94.3, -86.6) | **-65.9**<br>(-70.3, -61.7) |
| + 7 mM $\text{CaCl}_2$ (n=3) | **3.92**<br>(3.16, 4.84) | **-83.2**<br>(-85.2, -81.2) | **-52.3**<br>(-54.9, -49.8) |
| + 500 μM $\text{CaCl}_2$ (n=2) | **7.07**<br>(6.46, 7.70) | **-83.2**<br>(-84.3, -82.1) | **-53.4**<br>(-55.2, -52.4) |
| **$\text{CaCl}_2$** (n=2) | **ND** | **ND** | **ND** |
| + 400 μM **ATP** (n=4) | **58.9**<br>(49.4, 66.5) | **-11.3**<br>(-11.9, -10.1) | **12.8**<br>(11.9, 14.5) |
| **$\text{NaHCO}_3$** (n=2) | **ND** | **ND** | **ND** |

\* Binding parameters were determined by ITC at 25 °C in 20 mM HEPES, 150 mM NaCl, pH 8.0 using a single-site binding model. Global fitting was performed using data from replicate titrations (number of replicates indicated in table) to obtain best-fit values (bold). 68.3% confidence intervals for the fit of the model to the data are given in parentheses. ND, specific binding not detected.

**Table 2.** Data collection and refinement statistics for structures described in this work.

|  | apo-SbtB7001 | AMP:SbtB7001 | ADP:SbtB7001 | cAMP:SbtB7001 | ATP:SbtB7001 | Ca<sup>2+</sup>ATP:SbtB7001 |
| --- | --- | --- | --- | --- | --- | --- |
| PDB | 6N4A | 6MMO | 6MMC | 6MMQ | 6NTB | 6MM2 |
| **Data collection** |  |  |  |  |  |  |
| Space group | P 1 | P 1 | P 6<sub>3</sub> 2 2 | P 1 | P 6<sub>2</sub> | R 3 |
| Cell dimensions |  |  |  |  |  |  |
| *a*, *b*, *c* (Å) | 46.6 51.0 66.6 | 46.7 51.3 66.8 | 68.6 68.6 89.1 | 46.6 50.2 66.6 | 132.9 132.9 33.8 | 52.0 52.1 95.2 |
| $\alpha$, $\beta$, $\gamma$ (°) | 94.3 106.1 103.6 | 97.9 102.5 103.1 | 90 90 120 | 97.6 102.9 102.5 | 90 90 120 | 90 90 120 |
| Resolution (Å) | 49.03–2.30<br>(2.38–2.30) | 43.22–1.86<br>(1.93–1.86) | 49.44–1.42<br>(1.47–1.42) | 35.32–2.02<br>(2.09–2.02) | 43.52–1.90<br>(1.97–1.90) | 15.03–1.04<br>(1.08–1.04) |
| *R*<sub>merge</sub><br>(overall/inner/outer) | 0.040/0.015/1.05 | 0.046/0.033/1.922 | 0.105/0.037/6.445 | 0.05/0.029/1.270 | 0.107/0.054/2.680 | 0.039/0.035/1.337 |
| CC<sub>1/2</sub> | 0.999/0.999/0.574 | 0.998/0.996/0.335 | 1.000/1.000/0.380 | 0.998/0.998/0.307 | 0.999/0.999/0.560 | 0.999/0.997/0.405 |
| Completeness (%) | 98.0/96.6/95.2 | 97.1/86.5/96.3 | 100.0/99.7/100.0 | 96.4/93.5/96.6 | 100/99.3/100.0 | 95.2/53.4/50.9 |
| Multiplicity | 3.8/3.8/3.8 | 3.6/3.2/3.8 | 37.8/26.9/36.3 | 2.4/2.4/2.5 | 20/17.7/18.1 | 9.0/1.34/0.55 |
| **Refinement** |  |  |  |  |  |  |
| Resolution (Å) | 49.03–2.30<br>(2.38–2.30) | 41.58–1.86<br>(1.93–1.86) | 49.44–1.42<br>(1.47–1.42) | 35.32–2.02<br>(2.09–2.02) | 43.52–1.90<br>(1.97–1.90) | 15.03–1.04<br>(1.08–1.04) |
| No. of reflections | 24661 (2413) | 47132 (4612) | 24048 (2343) | 35629 (3583) | 27442 (2714) | 44104 (2995) |
| *R*<sub>work</sub> / *R*<sub>free</sub> | 0.2170/0.2767 | 0.2234/0.2579 | 0.2029/0.2202 | 0.2208/0.2794 | 0.210/0.2475 | 0.1452/1632 |
| No. of atoms | 3989 | 4189 | 788 | 4142 | 2541 | 997 |
| Protein | 3961 | 4002 | 722 | 3985 | 2350 | 845 |
| Ligands/Ions | 10 | 138 | 1 | 132 | 95 | 33 |
| Solvent | 18 | 49 | 65 | 34 | 96 | 119 |
| Average *B*-factor | 75.0 | 61.25 | 27.31 | 66.26 | 45.8 | 24.1 |
| Protein | 75.1 | 61.3 | 26.44 | 66.28 | 45.9 | 20.9 |
| Ligands/Ions | 84.12 | 59.9 | 28.82 | 60.20 | 38.8 | 15.5 |
| Solvent | 56.5 | 57.0 | 36.94 | 64.02 | 49.9 | 48.7 |
| R.m.s. deviations |  |  |  |  |  |  |
| Bond length (Å) | 0.011 | 0.007 | 0.005 | 0.008 | 0.007 | 0.013 |
| Bond angles (°) | 1.46 | 0.91 | 0.84 | 1 | 1.17 | 1.6 |

\*Statistics for the highest-resolution shell are shown in parentheses.

**Figure 1.**
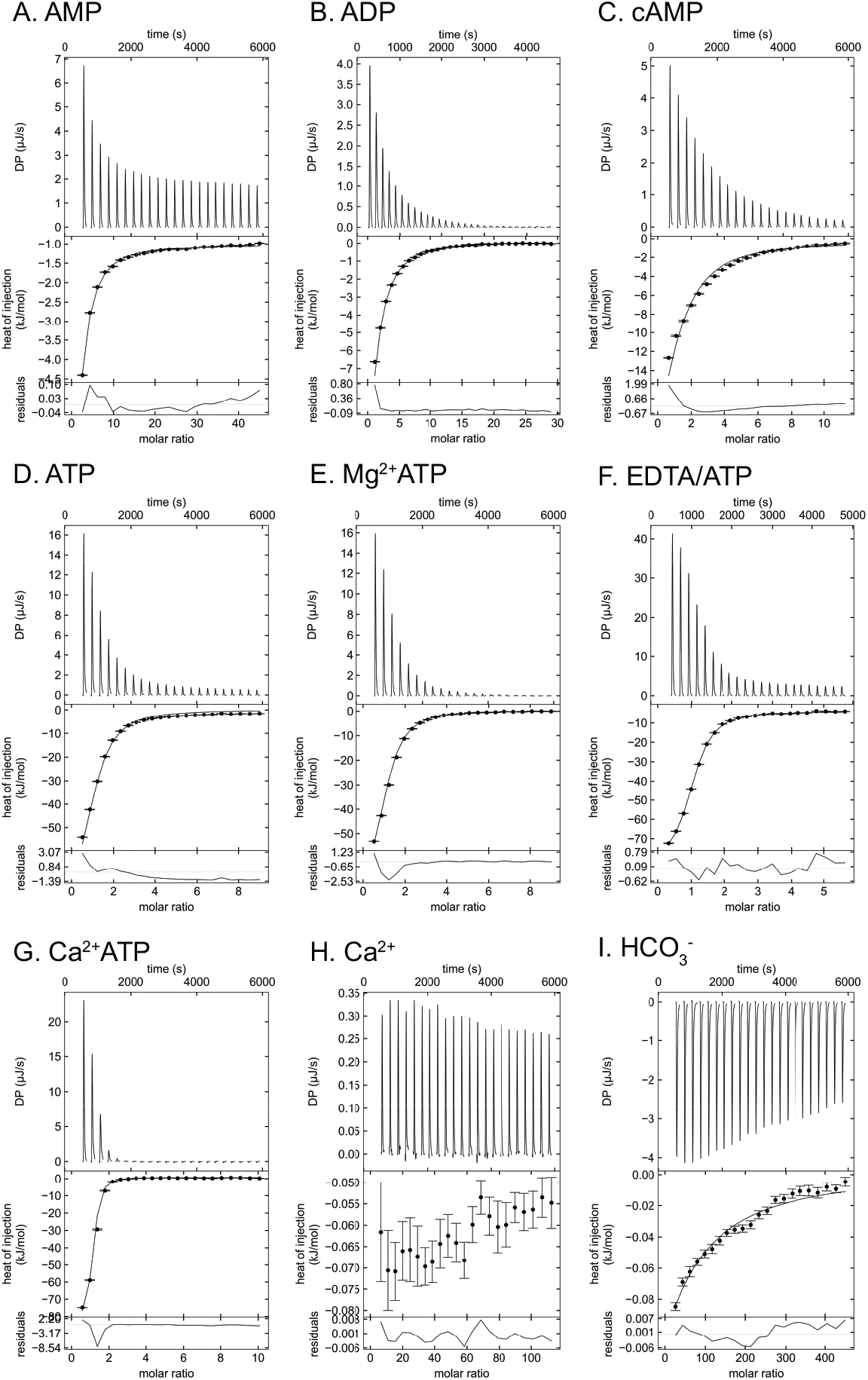
Isothermal titration calorimetry thermographs for the binding of nucleotides and potential effector molecules to SbtB7001. Representative ITC thermographs showing the titration of (A) 2 mM AMP into 110 μM SbtB7001, (B) 10 mM ADP into 110 μM SbtB7001, (C) 5 mM cAMP into 143 μM SbtB7001, (D) 4 mM ATP into 143 μM StbB7001, (E) 4 mM ATP into 143 μM SbtB7001 in the presence of 10 mM MgCl_2_, (F) 7 mM ATP into 400 μM SbtB7001 in the presence of 0.5 mM EDTA, (G) 4 mM ATP into 143 μM SbtB7001 in the presence of 7 mM CaCl_2_, (H) 50 mM CaCl_2_ into 143 μM SbtB7001 and (I) 200 mM NaHCO_3_ into 143 μM SbtB7001. The upper panels represent baseline-corrected power traces. The middle panels represent the integrated heat data and best fit (globally fitted data from replicate experiments where possible) of the independent binding site model in SEDPHAT [16]. Figures were produced using GUSSI [27]. The bottom panels show the residuals of the fit. Error bars represent the standard error in the integration of the peaks as calculated by NITPIC [15]. Additional ITC data are provided in **Supp. Figures 1-15**.

Since Mg^2+^ is often required for the binding of ATP to P_II_ proteins [12], we also tested the affinity of SbtB7001 for ATP in the presence of 10 mM Mg^2+^ (Figure 1E) and in the presence of the Mg^2+^-chelator EDTA (Figure 1F). The resulting values for the *K*_D_, ΔH, and TΔS of binding were not significantly different to when ATP was tested alone. We then tested the effect of Ca^2+^, which can also coordinate nucleotides [25,26]. ITC measurements revealed that the affinity for ATP was increased 10-fold, to ~4 μM (approximately 100-fold higher affinity than AMP in the absence of Ca^2+^) when 7 mM Ca^2+^ was present in the reaction conditions (Figure 1G). In the presence of a lower concentration of Ca^2+^ (500 μM), ATP binding was still enhanced, with a *K*_D_ of ~7 μM (Table 1, **Supp. Figure 11**). In the absence of nucleotides, binding of Ca^2+^ could not be detected by ITC (Figure 1H), suggesting that Ca^2+^ binding to SbtB7001 is dependent on pre-bound ATP or that Ca^2+^ and ATP bind cooperatively. Indeed, when Ca^2+^ was titrated into a solution of SbtB7001 in the presence of saturating concentrations of ATP (400 μM, > 10 × *K*_*D*_), isotherms indicated that Ca^2+^ bound with a *K*_D_ of approximately 60 μM (Table 1, **Supp. Figure 13**). The effect of Ca^2+^ on the binding of nucleotides was most pronounced for ATP; *K*_D_ values for cAMP and AMP were reduced by less than a factor of 2 and 4, respectively, in the presence of 500 μM Ca^2+^, and the *K*_D_ of ADP was unaffected by Ca^2+^ (Table 1, **Supp. Figure 4**).

Despite the role of SbtB in regulating SbtA-mediated HCO_3_^−^ transport, specific binding of HCO_3_^−^ to SbtB7001 could not be detected using ITC under the conditions tested (i.e. *K*_D_ for HCO_3_^−^ > 10 mM, Figure 1I). At the concentration of HCO_3_^−^ used in these titrations, the minor changes in heat observed in the control-subtracted thermograms are most likely due to non-specific binding events or artefacts. Indeed, when the same concentration of HCO_3_^−^ was titrated into the lac repressor (LacI) from *E. coli* similar heats were observed (**Supp. Figure 15**).

### 3.2 Crystal structures of SbtB7001

To understand the structural basis of nucleotide binding to SbtB7001, we crystallized and collected diffraction data on StbB7001 alone (P1, 2.3 Å) and co-crystallized SbtB7001 with AMP (P1, 1.86 Å), cAMP (P1, 2.02 Å), ADP (P6322, 1.42 Å), ATP (P62, 1.9 Å) and Ca^2+^ATP (R3, 1.04 Å). SbtB7001 is trimeric in solution (Figure 2A), and all crystal structures showed a trimeric P_II_-like core structure (Figure 2B) that was essentially identical to SbtB6803, with an RMSD of 0.45 Å between monomer α-carbons of apo-SbtB6803 (PDB: 5O3P) and apo-SbtB7001 (Figure 2C). The canonical nucleotide binding sites, which sit between the subunits of the trimer, are easily accessible from the solvent in apo-SbtB7001 (Figure 2D). StbB7001 shares 56% sequence identity with SbtB6803 (Figure 2E), which is consistent with the low RMSD value. However, SbtB6803 contains a C-terminal extension that is only present in a small subset of SbtB homologs, while SbtB7001 does not contain this extension.

**Figure 2.**
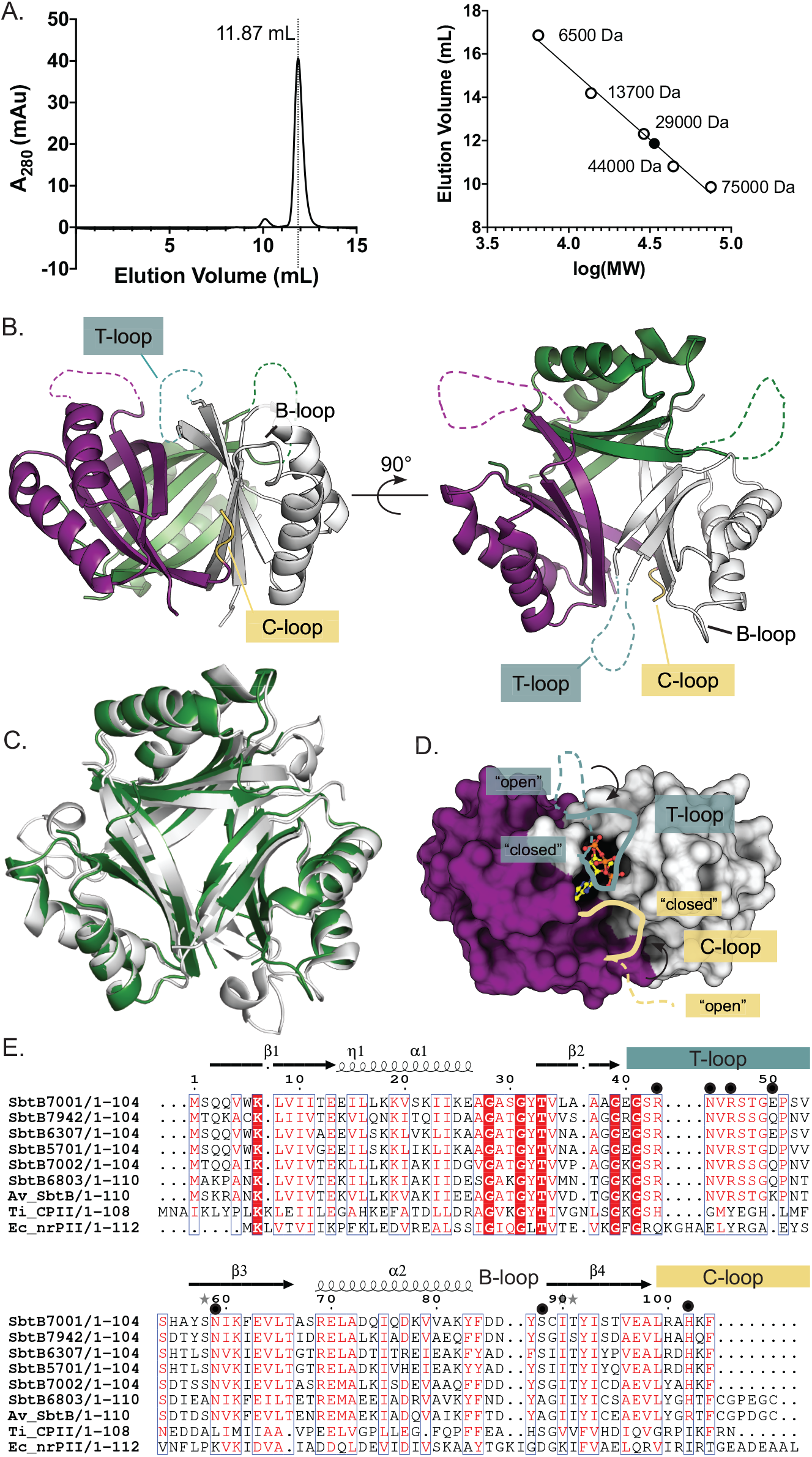
Overall fold of SbtB7001. (A) Size-exclusion chromatogram of SbtB7001 (left) and calibration curve for analytical size-exclusion chromatography (right). Open circles represent molecular weight standards and the closed circle represents untagged SbtB7001. The calculated molecular weight of SbtB7001 is consistent with a trimeric structure (calc. 33.7 kDa, theor. 34.5 kDa for the trimer). (B) Crystal structure of apo-SbtB7001 (Chains A, B, C; colored by chain) showing the conserved trimeric P_II_-like fold. The approximate positions of the flexible T-loop regions are indicated by the dashed lines, and the locations of the B-loop and C-loop are labelled. (C) Structural alignment of apo-SbtB7001 (green) and apo-SbtB6803 (PDB: 5O3P, white) showing conserved P_II_-like fold. (D) Nucleotides (e.g. ATP, shown as ball-and-stick) bind at the interface between subunits. The surface representation of SbtB7001 with T-loop and C-loop residues removed highlights the accessibility of the nucleotide binding site. Lines indicate the approximate position of “open” (dashed line) and “closed” (solid line) states of the T-loop and C-loop. (E) Sequence alignment of SbtB7001, SbtB homologs from other cyanobacterial strains and related P_II_ and P_II_-like proteins. Residues important for interactions with the phosphates of ATP are indicated by black circles. The primary and secondary structure of SbtB7001 is shown above the alignment. Sequences were aligned using Clustal Omega [28] and the figure was produced using EPScript 3.0 [29]. SbtB7001, *Cyanobium* sp. PCC7001 (WP_006910696.1); SbtB6803, *Synechocystis* sp. PCC6803 (WP_010872646.1); SbtB7942, *Synechococcus elongatus* PCC7942 (WP_011244772.1); SbtB6307, *Cyanobium* PCC6307 (WP_015110057.1; SbtB5701, *Synechococcus* WH5701 (WP_037981031.1); SbtB7002, *Synechococcus* sp. PCC7002 (WP_012306104.1); AvSbtB, *Anabaena variabilis* (yp_323533.1), TiCPII, carboxysome-associated P_II_ protein (CPII) from *Thiomonas Intermedia* K12 (WP_013104276.1); EcnrP_II_a, P_II_ protein from *Escherichia coli* (CCK45606.1).

### 3.3 Binding of AMP, ADP and cAMP at the inter-subunit clefts of SbtB7001

When SbtB7001 was co-crystallized with AMP (Figure 3A), ADP (Figure 3B), and cAMP (Figure 3C), electron density was observed for the respective ligands within each of the clefts between the monomers. The ADP:SbtB7001 co-crystal displayed ambiguous electron density around the β-phosphate of ADP, consistent with either a mobile β-phosphate, or hydrolysis of ADP and occupancy by AMP (as has been observed in other studies of related P_II_ proteins [10,11,30]).

**Figure 3.**
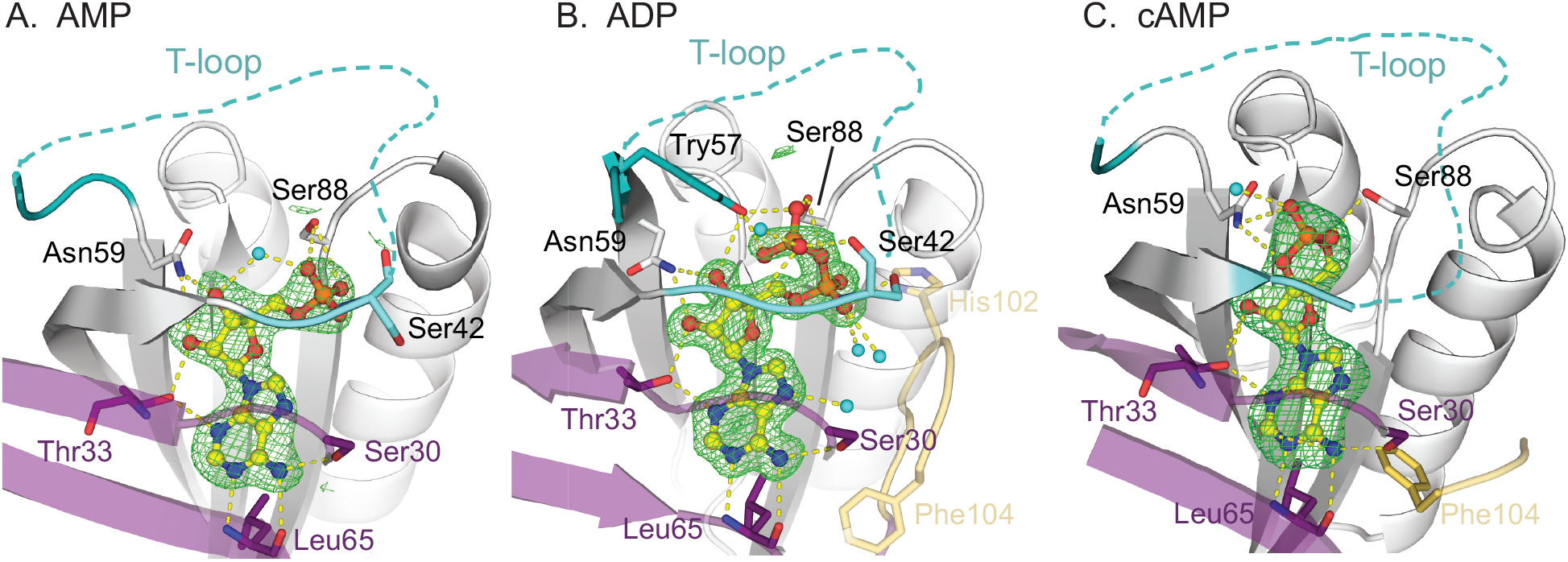
Ligand binding sites of SbtB7001. Representative binding sites from structures of SbtB7001 co-crystallized with (A) AMP, (B) ADP, and (C) cAMP. Ligands (yellow ball-and-stick) bind in the clefts between subunits (neighboring subunits are colored white and purple). Key residues for binding are shown as sticks. F_o_−F_c_ polder omit maps are shown as a green mesh contoured at 3σ and carved at 3 Å around the ligand. T-loop residues are colored cyan or teal, and the C-loop is colored yellow. A dashed line indicates the approximate location of the unmodelled, flexible T-loop. Coordinated water molecules are shown as cyan spheres.

Each nucleotide binds with the shared adenine motif sitting within a mostly hydrophobic cleft (lined by the side chains of Ile11, Leu65, Ile90), while the backbone carbonyl and amides of Gly31 and Leu65 and the hydroxyl groups of Ser30 and Thr33 form specific polar interactions with the base. The hydroxyl groups of the ribose sugar form hydrogen bonds with the side chains of Thr33 and Asn59 in each case, and the phosphate groups are positioned near the base of the flexible T-loop. In AMP:SbtB7001 (Figure 3A) and ADP:SbtB7001 (Figure 3B), the α-phosphate interacts with the side chain of Ser88, as well as with the backbone of Cys89 and Ser42. The interactions between the α-phosphate and the backbones of Ser42 and occasionally Gly41 seem to be important for stabilizing residues 40–42; in apo-SbtB7001 and cAMP:SbtB7001 these interactions are not observed, and this region becomes unstructured. Since AMP and ADP share common adenosine and ribose moieties that bind to SbtB7001 in an identical manner, the higher affinity of ADP (225 μM versus 387 μM; Table 1) must be attributed to differences in the way that the phosphate groups interact with SbtB7001. Indeed, in the ADP:Sbtb7001 complex we observe additional interactions between the β-phosphate and Ser88 and Tyr57, which is consistent with the ITC experiments that reveal a larger change in enthalpy upon binding of ADP compared with AMP (Table 1). Interactions between ADP and T-loop residues 40–56 are likely to only be transient, at most, since additional interactions with T-loop residues were not observed in the crystal structure of ADP:SbtB7001 and the electron density around the β-phosphate of ADP is consistent with a mobile β-phosphate that is not restrained by such interactions.

Due to the cyclic nature of cAMP, its binding mode is somewhat different to AMP/ADP; the interaction with Ser42 is not observed in cAMP:SbtB7001 and the phosphate forms an additional interaction with side chain of Asn59 (Figure 3C). Both SbtB7001 and SbtB6803 bind cAMP with higher affinity than AMP (e.g. 238 μM versus 447 μM for SbtB7001; Table 1) [10]. This is consistent with the crystal structures, which show the different position of the phosphate moiety of cAMP (versus AMP) facilitates an additional bond with Asn59, a residue that is conserved amongst SbtB homologs (Figure 1E). This interaction is not observed in AMP:SbtB7001 and could explain the increased affinity for cAMP by SbtB7001 and SbtB6803 [10].

### 3.4 Ca^2+^ and ATP stabilize the T-loop

In contrast to AMP and ADP, the γ-phosphate of ATP supports additional interactions with the highly conserved residues at the base of the T-loop (residues 40–46) (Figure 4A, 4B), explaining why SbtB7001 binds ATP with a much higher affinity and has a more favorable enthalpy of binding compared with AMP, ADP or cAMP. The salt bridge formed between the γ-phosphate of ATP and the side chain of Arg46 appears to be particularly important, given the high conservation of this residue among SbtB homologs (Figure 1E). This ATP-specific interaction appears to stabilize an interaction between Arg46 and Asn59, which pulls the base of the T-loop towards the core of the protein, where it tightly encircles the phosphate groups of the ligand. In canonical P_II_-proteins, the residue at the position analogous to Asn59 in SbtB7001 is involved in forming a key salt bridge with T-loop residues, and the formation and breaking of this salt bridge has been shown to play an important role in conferring functionally-relevant T-loop conformations [31,32]. In SbtB7001, when the T-loop is held in position by the Arg46–Asn59 interaction, additional interactions are also formed between the phosphates of ATP, Ser88, Arg43 and multiple points on the backbone of the T-loop, which further stabilize this region. While the interaction between the α-phosphate and Arg43 could be possible when AMP or ADP is bound, it seems that the additional ATP-specific interaction with Arg46 is required to first stabilize this basal T-loop region and promote this interaction between the α-phosphate and Arg43. Thus, in comparison to the largely disordered T-loop residues 43–55 in apo-SbtB7001, AMP:SbtB7001, ADP:SbtB7001 and cAMP:SbtB7001 (as was also observed in structures of SbtB6803 bound to AMP and cAMP [10]), the binding of ATP to SbtB7001 resulted in stabilization of the T-loop. These structural observations are consistent with our ITC experiments for ATP, which revealed a substantially greater entropic cost associated with ATP binding, compared with the binding of AMP, ADP and cAMP (Table 1).

**Figure 4.**
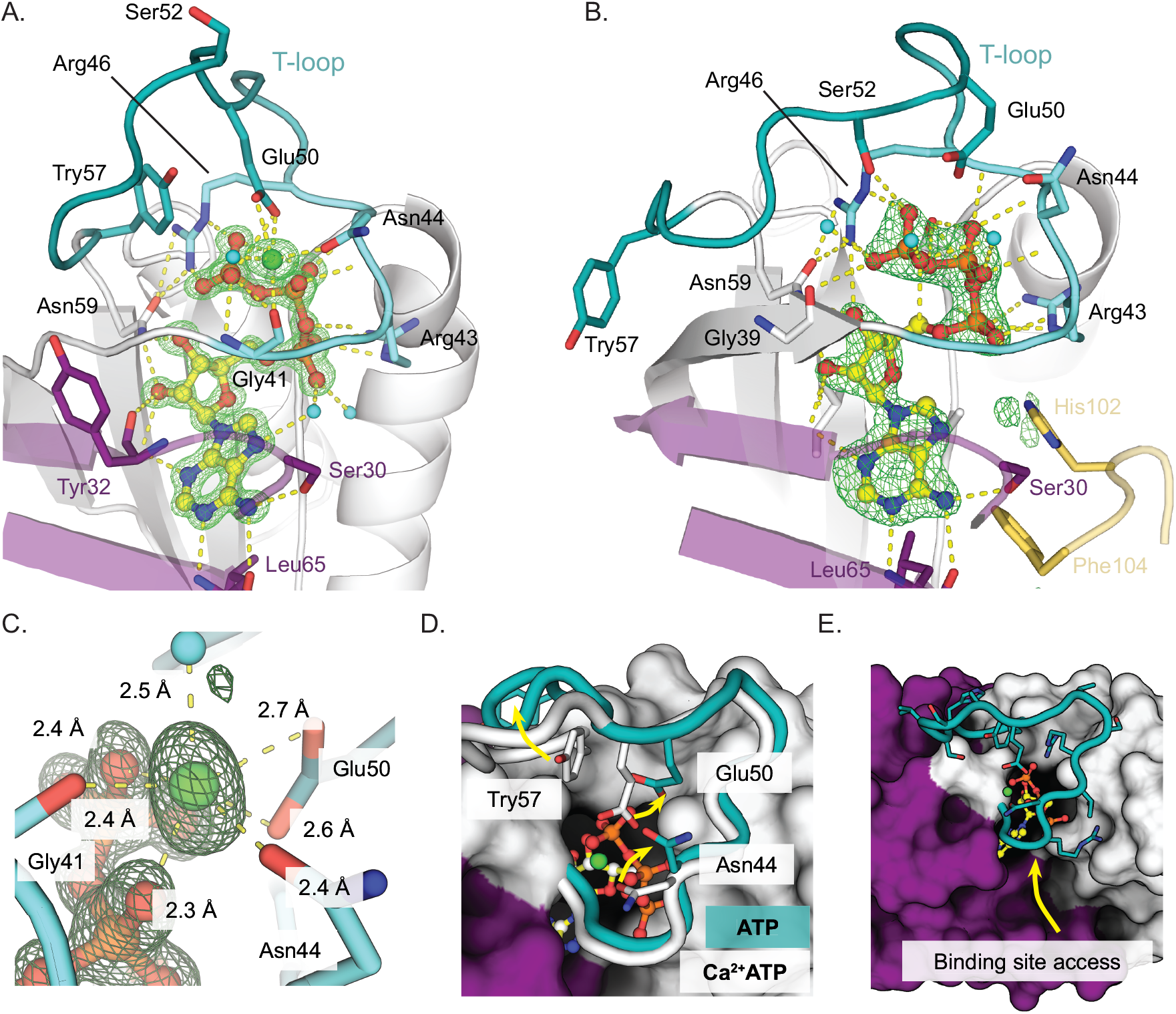
Ca^2+^ATP and ATP stabilize the T-loop of SbtB7001. (A) Ligand binding site of Ca^2+^ATP:SbtB7001. Ligands are shown as yellow ball-and-sticks and key residues for binding are shown as sticks. F_o_−F_c_ polder omit maps are shown as a green mesh contoured at 3σ and carved at 3 Å around the ligand. T-loop residues are colored cyan (residues 40–46) or teal (residues 47–57). The C-loop is colored yellow. Coordinated water molecules are shown as cyan spheres. (B) Ligand binding site of ATP:SbtB7001. (C) Close up of Ca^2+^ binding site showing distances to coordinating atoms. (D) When completely modelled, the T-loop of ATP:SbtB7001 (teal) forms a conformation distinct from that observed in Ca^2+^ATP (white). Movement of key residues is indicated by yellow arrows. (E) The T-loop is stabilized against the core of the protein by Ca^2+^ATP: and occludes the binding site from above. The binding site can still be accessed from the direction of the yellow arrow.

Although both ATP:SbtB7001 and Ca^2+^ATP:StbB7001 co-crystals were grown in the presence of 10 mM Mg^2+^, electron density consistent with an octahedrally coordinated metal ion near the γ-phosphate of ATP was only observed in the crystal grown in the presence of 20 mM Ca^2+^ (Figure 4B). Metal−oxygen distances (~2.4 Å) and environmental B-factor analysis (B-factor 20.38 for Ca^2+^, averaged B-factor of 21.94 for coordinated atoms) are consistent with a Ca^2+^ binding site. This was confirmed using the ‘place elemental ions’ tool in *phenix.refine* (valence sum for Ca^2+^ is 1.989, expected 2; valence sum for Mg^2+^ is 0.948, expected 2) [33]. The Ca^2+^ ion is coordinated by the side chains of Glu50 and Asn44, the backbone carbonyl of Gly41, the β- and γ-phosphates of ATP, and a single water molecule that is held in place by Ser54 (Figure 4C). The interactions with Glu50 and Asn44 appear to be important for stabilizing the apex of the T-loop; when Ca^2+^ was absent from the crystallization conditions, the complete T-loop could only be completely modelled in one of the three chains (ATP:SbtB7001, chain C), consistent with less order between residues 46 and 57. Even when this region could be completely modelled, the T-loop of ATP:SbtB7001 adopts a conformation distinct from that seen in Ca^2+^ATP:SbtB7001 (Figure 4D): Asn44 and Tyr57 flip away from the Ca^2+^ binding site, Glu50 moves away from the ATP binding site, and Pro51 adopts a cis-conformation. Thus, it seems that these Ca^2+^-specific interactions further stabilize the ‘closed’ conformation of the T-loop, which is consistent with our observation from ITC that affinity for ATP was increased in the presence of Ca^2+^, but not Mg^2+^. This ‘closed’ Ca^2+^ATP:SbtB7001 T-loop conformation almost completely seals the nucleotide binding site from one side, leaving only a narrow channel near the β- and γ-phosphates that could allow for the passage of an ion to and from the metal binding site, as well a thin access tunnel near the C-terminus of the protein (Figure 4E). In contrast, the flexibility observed in the ATP:SbtB7001 T-loop in the absence of Ca^2+^ means that the phosphate tail of ATP is not completely occluded by the T-loop.

### 3.5 The dynamic C-loop forms part of the nucleotide binding site

The C-terminal C-loop forms part of the nucleotide binding site in canonical P_II_ proteins [34]. In SbtB7001, the C-terminal residues Leu99–Phe104 could only be completely modelled in AMP:SbtB7001 (Chain A only), cAMP:SbtB7001 (Chain B only), ADP:SbtB7001, and ATP:SbtB7001 (Figure 5). All other chains showed poor electron density in this region, consistent with conformational heterogeneity. In crystals where the C-terminal residues could be modelled, this region occupies part of the inter-subunit cleft, where it forms part of the ligand-binding cavity, occludes access to the binding site, and can come within bonding distance of both the ligand and the T-loop.

**Figure 5.**
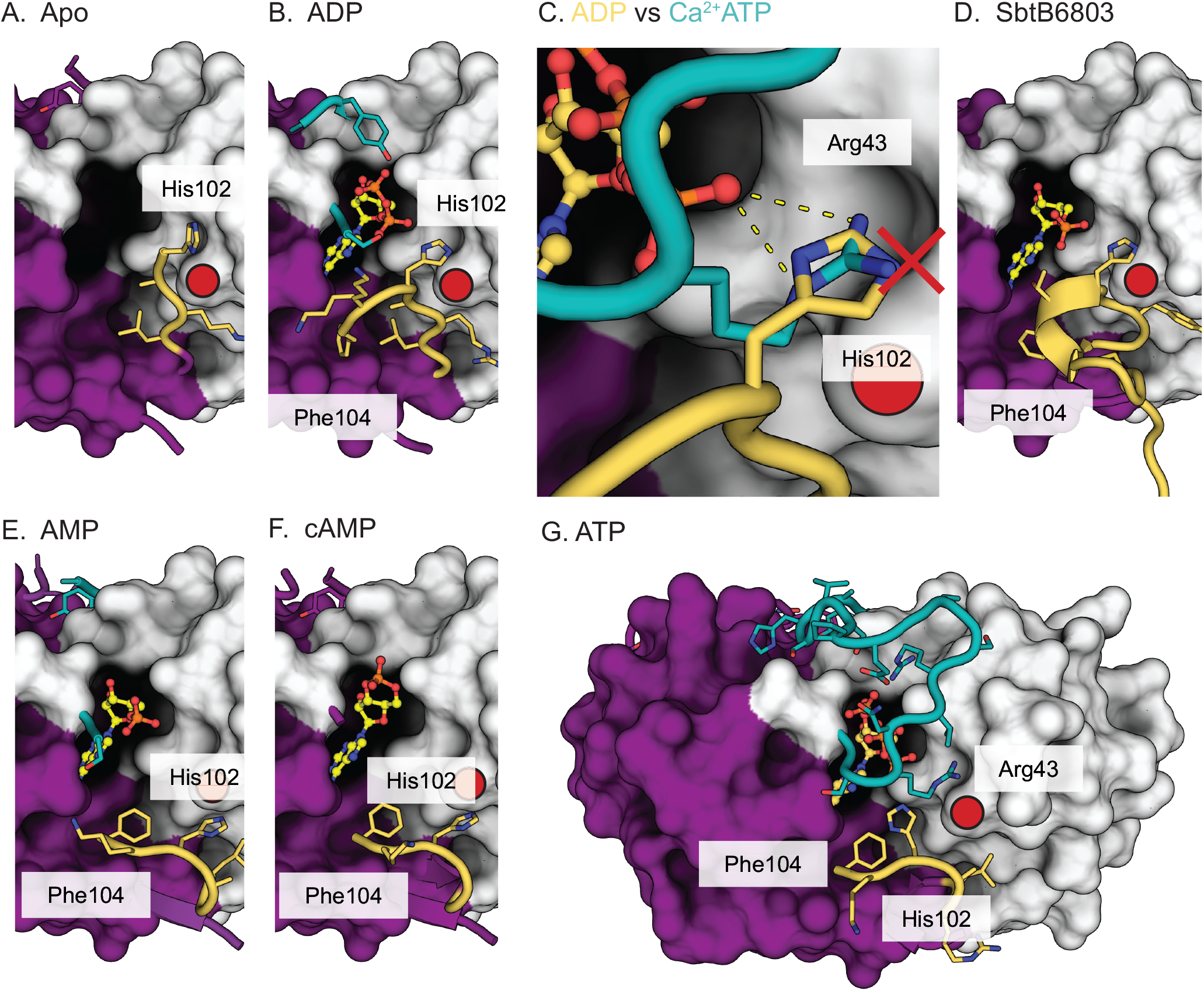
Structural changes at C-loop of SbtB7001. The C-loop (residues 99–104, yellow) adopted distinct conformations in the crystal structures of SbtB7001 and SbtB6803. T-loop residues are colored teal. The position of Phe88 is indicated by the red circle. In apo-SbtB7001 (A) and ADP:SbtB7001 (B) the C-loop extends up into the intersubunit cleft. These conformations would not be compatible with the closed T-loop observed when ATP is bound; for example, in ADP:SbtB7001 the C-loop residue His102 occupies the space that T-loop residue Arg43 occupies in Ca^2+^ATP:SbtB7001 (C). The rigid C-terminal region of SbtB6803 (e.g. PDB: 5O3R) holds His102 in a similar position that would likely clash with the closed T-loop. (E-G) The C-loop adopts a more compact pose in chains of AMP:SbtB7001 (E), cAMP:SbtB7001 (F) and ATP:SbtB7001 (G). As exemplified in ATP:SbtB7001, this conformation is compatible with the closed T-loop conformation. When the C-loop and T-loop are both closed, the nucleotide binding site is completely occluded.

In apo-SbtB7001, residues Leu99–His102 adopt an extended conformation: these residues run antiparallel to β4 of the neighboring subunit with backbone hydrogen bonds holding it in place (Figure 5A). In this pose, His102 is positioned between Phe84 and the nucleotide binding site. His102 is located in a similar position in ADP:SbtB7001, where it is within hydrogen bonding distance (~3 Å) of the α-phosphate of the ligand (Figure 5B). While this interaction could aid in binding the α-phosphate of ADP/AMP, it is sterically incompatible with the ‘closed’ position of the T-loop observed in Ca^2+^ATP:SbtB7001 and ATP:SbtB7001, clashing with the position of Arg43 (Figure 5C). The rigid C-terminal helix observed in SbtB6803 holds His102 in a similar location and would cause a similar steric clash with the position of Arg43 that appears to be important for ATP-binding (Figure 5D).

In AMP:SbtB7001 (chain A, Figure 5E) and cAMP:SbtB7001 (chain B, Figure 5F), β4 continues to run antiparallel with β1 until Ala101 before wrapping back around towards the ligand-binding site. In this more compact conformation, His102 is no longer within bonding distance of the ligand. The C-loop of ATP:SbtB7001 traces a similar route, although His102 adopts an alternate rotamer that positions it within bonding distance of the α-phosphate of ligand (Figure 5G). Unlike the extended conformation observed in apo-SbtB7001 and ADP:SbtB7001, the more compact conformation observed in AMP:SbtB7001, cAMP:SbtB7001 and ATP:SbtB7001 is compatible with the closed conformation of the T-loop (e.g. as observed in ATP:SbtB7001). When both the T-loop and C-loop adopt a closed conformation (as in ATP:SbtB7001), the nucleotide binding site access is almost completely occluded.

In the related, carboxysome-associated P_II_ protein from *Thiomonas intermedia* (TiCPII), HCO_3_^−^ binds near the C-loop and alters the affinity of the protein towards nucleotides [11]. While most ligand-bound crystals of SbtB7001 were grown in the presence of 50–100 mM NaHCO_3_, electron density that unambiguously corresponded to HCO_3_^−^ could not be identified in any of the structures. This is consistent with our ITC results that showed only low-affinity, binding of HCO_3_^−^ to SbtB7001, consistent with non-specific protein stabilization.

## 4. DISCUSSION

In this work we have shown that, like SbtB6803 [10], SbtB7001 binds AMP, ADP, cAMP and ATP with micromolar-range affinities. However, in contrast to SbtB6803, we found that SbtB7001 binds ATP with an affinity 5-to 10-fold greater than the other nucleotides tested, and that the affinity for ATP increases a further 10-fold (from a *K*_D_ of ~40 μM to ~4 μM) in the presence of Ca^2+^. In contrast, the presence of Ca^2+^ had a less substantial effect on the binding of AMP, ADP or cAMP. ITC experiments revealed that binding of ATP and Ca^2+^ATP to SbtB7001 is associated with a large change in enthalpy and loss of entropy, consistent with the formation of a number of charged interactions between the ligand and SbtB7001, and a significant stabilization of a mobile region of the protein, respectively. Structures of SbtB7001 co-crystallized with ligands support the ITC results and (i) provide a structural explanation for the preferential binding of ATP, (ii) account for differences in the affinities of SbtB7001 and those reported for SbtB6803 [10], and (iii) reveal how binding of ATP stabilizes the otherwise flexible T-loop. Together, these results provide a structural explanation for how SbtB could regulate SbtA, via changes in the adenylate charge ratio ([ATP]/[ATP+ADP+AMP]) observed in a transition from light to darkness in cyanobacteria [35].

The mode of ATP binding to SbtB7001 is distinct from ATP binding in the canonical P_II_ proteins, in which a conserved RXR motif in the C-terminal C-loop is responsible for forming interactions with the terminal phosphates of the ligand [34]. In SbtB7001 these interactions are instead provided by arginine residues from the T-loop. Indeed, in SbtB7001 (and the majority of the cyanobacterial SbtBs; Figure 1E) the C-terminal extension is largely truncated, suggesting that it is less essential for ligand binding in SbtBs from most cyanobacteria. The reduced C-loop in SbtB7001 displayed conformational heterogeneity, adopting extended and compact states, with the extended conformation being incompatible with Ca^2+^ATP binding. It is unlikely that the C-loop substantially contributes to the differentiation between AMP, ADP, or ATP, since C-loop residues (e.g. His102) are unlikely to be able to form interactions with the β- or γ-phosphates of ADP or ATP.

The T-loop residues involved in binding ATP are highly conserved amongst most SbtB homologs (Figure 1E), suggesting that all SbtB homologs should bind ATP more tightly than AMP, ADP or cAMP. However, this was not observed for Sbtb6803 [10]. The C-terminal extension found in SbtB6803, which is only present in a small subset of SbtB homologs, may contribute to the lower affinity between SbtB6803 and ATP. Indeed, based on the structures presented here, the rigid, disulfide-linked C-terminal region of SbtB6803 would clash with the position that Arg43 is required to adopt for the T-loop to accommodate ATP. Interestingly, the C-terminal region is unresolved in the structure of a related SbtB from *Anabaena Variabilis* ATCC 29413 (AvSbtB, PDB: 3DFE), which also has the putative redox-sensing C-terminal extension, suggesting that this region is able to switch between open and closed states. It is therefore possible that that the C-terminal extension found in SbtB6803 and a small subset of SbtBs could alter the affinity of SbtB for ATP based on the cellular redox conditions of the cell by preventing conformations of the T-loop that are required for ATP binding [10]. The ATP-binding activity of SbtB6803 was not tested in different reducing/oxidizing conditions [10], and it is possible that SbtB6803 has a higher affinity for ATP under certain redox conditions.

In the cyanobacterial cell, the ligand-bound state of SbtB will depend on both the affinity of the ligands, and the local concentrations of each nucleotide and/or effector molecules. Our results suggest that the apo-form of SbtB7001 is unlikely to be physiologically relevant because there will always be AMP/ADP/cAMP/ATP present at concentrations in excess of the *K*_D_ [36]. The ratio of [ATP]/[ATP+ADP] has been reported to be around 0.8 in *Synechococcus* under illumination [37], but varies significantly in cyanobacteria in response to changing light and C_i_ availability. For example, the [ATP]/[ADP+ATP] ratio drops from 0.9 to around 0.4 within 5-6 hours of *Synechococcus elongatus* PCC7942 being moved to darkness, and returns to about 0.85 within an hour of being returned to the light [35]. Thus, when ATP levels drop (such as during darkness), the binding sites of SbtB7001 are likely to become occupied by ADP (or possibly by AMP or cAMP), rather than ATP. Since the SbtA-SbtB6803 complex is stabilized by AMP and ADP [10], we expect that binding of these ligands would also stabilize the inactive SbtA-SbtB7001 complex (with open T-loops) to prevent futile metabolic cycling in the dark. We envision that when ATP levels rise again (upon exposure to light), ATP could displace AMP/ADP and cause the T-loops to adopt the closed conformation, which we envisage would be the allosteric signal to cause the complex to dissociate. Further *in vivo* studies will be required to test this model.

The finding that Ca^2+^ enhances ATP binding to SbtB7001 raises the possibility that Ca^2+^ might act a second messenger to influence how SbtB regulates SbtA-mediated transport. Interestingly, transient increases in Ca^2+^ have been shown to upregulate expression of CCM-related genes, including the *sbt* operon, in *Anabaena* sp. PCC7120 [38]. Ca^2+^ is known to play an important role in the regulation of other cyanobacterial systems as well. For example, Ca^2+^ acts as a signal for heterocyst formation [39] and other regulatory proteins involved in sensing C_i_, such as soluble adenylyl cyclases, can simultaneously sense ATP, Ca^2+^, and HCO_3_^−^/CO_2_/pH [40]. Further evidence that Ca^2+^ plays an important role in the regulation of cyanobacterial HCO_3_^−^ transport includes a structure of the periplasmic domain of another cyanobacterial HCO_3_^−^ transporter, CmpA, which revealed that HCO_3_^−^ bound strongly only if Ca^2+^ was present [41]. Future experiments could focus on testing whether physiological levels of Ca^2+^ could act as a secondary effector for SbtB *in vivo* since free calcium levels in cyanobacteria can vary between 100–200 nM in normal conditions [42], to low micromolar-range concentrations following light to dark transitions [43] or heat-, cold-, or ammonia-shock [44]. If ATP binding does cause the complex to dissociate, then variations in the level of Ca^2+^ in the cell could further control this equilibrium by altering the affinity of SbtB for ATP. For example, transient increases in the concentration of free Ca^2+^ following light-to-dark transitions could conceivably help delay the formation of the SbtA-SbtB complex following the onset of darkness.

In summary, this study has provided a molecular basis for the design of future *in vivo* studies to test a number of hypotheses relating to the regulation of the SbtA-SbtB complex that will be required to propose a comprehensive model describing the regulation of SbtA-mediated HCO_3_^−^ transport by SbtB. First, rising ATP concentrations (due to exposure to light) are likely to be an allosteric signal to release the inhibitory SbtB protein from the StbA HCO_3_^−^ transporter. Second, fluctuations in Ca^2+^ levels are likely to modulate the effect of changing nucleotide concentrations. Third, the C-terminal extension, which is present in a subset of SbtBs, may play an additional role in modulating allosteric signaling by controlling ATP binding in a redox-sensitive fashion. These findings will guide future experiments that will ultimately contribute to attempts to use transgenic SbtA to improve the photosynthetic performance and yield of C3 crop plants by increasing the steady state concentration of C_i_ in the chloroplast stroma [2].

## ABBREVIATIONS (non-standard)

C_i_: inorganic carbon
CCM: CO_2_-concentrating mechanism
SbtA: sodium-dependent bicarbonate transporter A
SbtB: sodium-dependent bicarbonate transport protein B

## Funding

This work was supported by the Australian Government through the Australian Research Council Centre of Excellence for Translational Photosynthesis (CE1401000015). JAK acknowledges support from an Australian Government Research Training Program Scholarship.

We would like to thank staff at the Collaborative Crystallisation Centre (C3, CSIRO) for assistance in crystallizing these proteins. We thank Li Lynn Tan (ANU) and Australian Synchrotron beamline scientists for help with freezing crystals and the collection of diffraction data.

## ACCESSION CODES

The atomic coordinates and structure factors for apo-SbtB7001 (PDB: 6N4A), AMP:SbtB7001 (PDB: 6MMO), ADP:SbtB7001 (PDB: 6MMC), cAMP:SbtB7001 (PDB: 6MMQ), ATP:SbtB7001 (PDB: 6NTB) and Ca^2+^ATP (PDB: 6MM2) have been deposited in the Protein Data Bank (http://wwpdb.org/).

## REFERENCES

[1] G.D. Price, M.R. Badger, F.J. Woodger, B.M. Long, Advances in understanding the cyanobacterial CO_2_-concentrating-mechanism (CCM): functional components, Ci transporters, diversity, genetic regulation and prospects for engineering into plants, J. Exp. Bot. 59, 7 (2008) 1441–1461.

[2] B.M. Long, B.D. Rae, V. Rolland, B. Forster, G.D. Price, Cyanobacterial CO_2_-concentrating mechanism components: function and prospects for plant metabolic engineering, Curr. Opin. Plant Biol. 31, (2016) 1–8. 10.1016/j.pbi.2016.03.002

[3] G.D. Price, Inorganic carbon transporters of the cyanobacterial CO_2_ concentrating mechanism, Photosynth. Res. 109, 1–3 (2011) 47–57. 10.1007/s11120-010-9608-y

[4] R.L. Burnap, M. Hagemann, A. Kaplan, Regulation of CO_2_ Concentrating Mechanism in Cyanobacteria, Life (Basel) 5, 1 (2015) 348–371. 10.3390/life5010348

[5] H.L. Wang, B.L. Postier, R.L. Burnap, Alterations in global patterns of gene expression in *Synechocystis* sp. PCC 6803 in response to inorganic carbon limitation and the inactivation of ndhR, a LysR family regulator, J. Biol. Chem. 279, 7 (2004) 5739–5751. 10.1074/jbc.M311336200

[6] F.J. Woodger, D.A. Bryant, G.D. Price, Transcriptional regulation of the CO_2_-concentrating mechanism in a euryhaline, coastal marine cyanobacterium, *Synechococcus* sp. strain PCC 7002: role of NdhR/CcmR, J. Bacteriol. 189, 9 (2007) 3335–3347. 10.1128/JB.01745-06

[7] M. Eisenhut, E. Aguirre von Wobeser, L. Jonas, H. Schubert, B.W. Ibelings, H. Bauwe, H.C. Matthijs, M. Hagemann, Long-term response toward inorganic carbon limitation in wild type and glycolate turnover mutants of the cyanobacterium *Synechocystis* sp. strain PCC 6803, Plant Physiol. 144, 4 (2007) 1946–1959. 10.1104/pp.107.103341

[8] M. Shibata, H. Katoh, M. Sonoda, H. Ohkawa, M. Shimoyama, H. Fukuzawa, A. Kaplan, T. Ogawa, Genes essential to sodium-dependent bicarbonate transport in cyanobacteria: function and phylogenetic analysis, J. Biol. Chem. 277, 21 (2002) 18658–18664. 10.1074/jbc.M112468200

[9] J.H. Du, B. Forster, L. Rourke, S.M. Howitt, G.D. Price, Characterisation of cyanobacterial bicarbonate transporters in *E. coli* shows that SbtA homologs are functional in this heterologous expression system, PLoS ONE 9, 12 (2014) e115905. 10.1371/journal.pone.0115905

[10] K.A. Selim, F. Haase, M.D. Hartmann, M. Hagemann, K. Forchhammer, PII-like signaling protein SbtB links cAMP sensing with cyanobacterial inorganic carbon response, Proc. Natl. Acad. Sci. U. S. A. 115, 21 (2018) E4861–E4869. 10.1073/pnas.1803790115

[11] N.M. Wheatley, K.D. Eden, J. Ngo, J.S. Rosinski, M.R. Sawaya, D. Cascio, M. Collazo, H. Hoveida, W.L. Hubbell, T.O. Yeates, A PII-like protein regulated by bicarbonate: Structural and biochemical studies of the carboxysome-associated CPII protein, J. Mol. Biol. 428, 20 (2016) 4013–4030. 10.1016/j.jmb.2016.07.015

[12] K. Forchhammer, J. Luddecke, Sensory properties of the PII signalling protein family, FEBS J. 283, 3 (2016) 425–437. 10.1111/febs.13584

[13] M. Radchenko, M. Merrick, The role of effector molecules in signal transduction by PII proteins, Biochem. Soc. Trans. 39, 1 (2011) 189–194. 10.1042/BST0390189

[14] A.M. Catanzariti, T.A. Soboleva, D.A. Jans, P.G. Board, R.T. Baker, An efficient system for high-level expression and easy purification of authentic recombinant proteins, Protein Sci. 13, 5 (2004) 1331–1339. 10.1110/ps.04618904

[15] S. Keller, C. Vargas, H. Zhao, G. Piszczek, C.A. Brautigam, P. Schuck, High-precision isothermal titration calorimetry with automated peak-shape analysis, Anal. Chem. 84, 11 (2012) 5066–5073. 10.1021/ac3007522

[16] H. Zhao, G. Piszczek, P. Schuck, SEDPHAT--a platform for global ITC analysis and global multi-method analysis of molecular interactions, Methods 76, (2015) 137–148. 10.1016/j.ymeth.2014.11.012

[17] W. Kabsch, Xds, Acta Crystallogr. D Biol. Crystallogr. 66, Pt 2 (2010) 125–132. 10.1107/S0907444909047337

[18] P.R. Evans, G.N. Murshudov, How good are my data and what is the resolution?, Acta Crystallogr. D Biol. Crystallogr. 69, Pt 7 (2013) 1204–1214. 10.1107/S0907444913000061

[19] A. Vagin, A. Teplyakov, MOLREP: an automated program for molecular replacement, J. Appl. Crystallogr. 30, (1997) 1022–1025. Doi 10.1107/S0021889897006766

[20] A.J. McCoy, R.W. Grosse-Kunstleve, P.D. Adams, M.D. Winn, L.C. Storoni, R.J. Read, Phaser crystallographic software, J. Appl. Crystallogr. 40, Pt 4 (2007) 658–674. 10.1107/S0021889807021206

[21] P. Emsley, K. Cowtan, Coot: model-building tools for molecular graphics, Acta Crystallogr. D Biol. Crystallogr. 60, Pt 12 Pt 1 (2004) 2126–2132. 10.1107/S0907444904019158

[22] G.N. Murshudov, A.A. Vagin, E.J. Dodson, Refinement of macromolecular structures by the maximum-likelihood method, Acta Crystallogr. D Biol. Crystallogr. 53, Pt 3 (1997) 240–255. 10.1107/S0907444996012255

[23] P.V. Afonine, R.W. Grosse-Kunstleve, N. Echols, J.J. Headd, N.W. Moriarty, M. Mustyakimov, T.C. Terwilliger, A. Urzhumtsev, P.H. Zwart, P.D. Adams, Towards automated crystallographic structure refinement with *phenix.refine*, Acta Crystallogr. D Biol. Crystallogr. 68, Pt 4 (2012) 352–367. 10.1107/S0907444912001308

[24] R.P. Joosten, F. Long, G.N. Murshudov, A. Perrakis, The PDB_REDO server for macromolecular structure model optimization, IUCrJ 1, Pt 4 (2014) 213–220. 10.1107/S2052252514009324

[25] M.S. Mohan, G.A. Rechnitz, Ion-electrode study of the calcium-adenosine triphosphate system, J. Am. Chem. Soc. 94, 5 (1972) 1714–1716. 10.1021/ja00760a048

[26] M. Picard, A.M. Jensen, T.L. Sorensen, P. Champeil, J.V. Moller, P. Nissen, Ca^2+^ versus Mg^2+^ coordination at the nucleotide-binding site of the sarcoplasmic reticulum Ca^2+^-ATPase, J. Mol. Biol. 368, 1 (2007) 1–7. 10.1016/j.jmb.2007.01.082

[27] C.A. Brautigam, Calculations and publication-quality illustrations for analytical ultracentrifugation data, Methods Enzymol. 562, (2015) 109–133. 10.1016/bs.mie.2015.05.001

[28] F. Sievers, D.G. Higgins, Clustal Omega, accurate alignment of very large numbers of sequences, Methods Mol. Biol. 1079, (2014) 105–116. 10.1007/978-1-62703-646-7_6

[29] X. Robert, P. Gouet, Deciphering key features in protein structures with the new ENDscript server, Nucleic Acids Res. 42, Web Server issue (2014) W320–324. 10.1093/nar/gku316

[30] M.V. Radchenko, J. Thornton, M. Merrick, PII signal transduction proteins are ATPases whose activity is regulated by 2-oxoglutarate, Proc. Natl. Acad. Sci. U. S. A. 110, 32 (2013) 12948–12953. 10.1073/pnas.1304386110

[31] S. Maier, P. Schleberger, W. Lu, T. Wacker, T. Pfluger, C. Litz, S.L. Andrade, Mechanism of disruption of the Amt-GlnK complex by PII-mediated sensing of 2-oxoglutarate, PLoS ONE 6, 10 (2011) e26327. 10.1371/journal.pone.0026327

[32] M.V. Radchenko, J. Thornton, M. Merrick, Control of AmtB-GlnK complex formation by intracellular levels of ATP, ADP, and 2-oxoglutarate, J. Biol. Chem. 285, 40 (2010) 31037–31045. 10.1074/jbc.M110.153908

[33] N. Echols, N. Morshed, P.V. Afonine, A.J. McCoy, M.D. Miller, R.J. Read, J.S. Richardson, T.C. Terwilliger, P.D. Adams, Automated identification of elemental ions in macromolecular crystal structures, Acta Crystallogr. D Biol. Crystallogr. 70, Pt 4 (2014) 1104–1114. 10.1107/S1399004714001308

[34] K. Forchhammer, PII signal transducers: novel functional and structural insights, Trends Microbiol. 16, 2 (2008) 65–72. 10.1016/j.tim.2007.11.004

[35] M.J. Rust, S.S. Golden, E.K. O’Shea, Light-driven changes in energy metabolism directly entrain the cyanobacterial circadian oscillator, Science 331, 6014 (2011) 220–223. 10.1126/science.1197243

[36] O. Fokina, C. Herrmann, K. Forchhammer, Signal-transduction protein PII from *Synechococcus elongatus* PCC 7942 senses low adenylate energy charge in vitro, Biochem. J. 440, 1 (2011) 147–156. 10.1042/BJ20110536

[37] T. Kallas, R.W. Castenholz, Internal pH and ATP-ADP pools in the cyanobacterium *Synechococcus* sp. during exposure to growth-inhibiting low pH, J. Bacteriol. 149, 1 (1982) 229–236.

[38] J. Walter, F. Lynch, N. Battchikova, E.M. Aro, P.J. Gollan, Calcium impacts carbon and nitrogen balance in the filamentous cyanobacterium *Anabaena* sp. PCC 7120, J. Exp. Bot. 67, 13 (2016) 3997–4008. 10.1093/jxb/erw112

[39] Y.H. Zhao, Y.M. Shi, W.X. Zhao, X. Huang, D.H. Wang, N. Brown, J. Brand, J.D. Zhao, CcbP, a calcium-binding protein from *Anabaena* sp PCC7120 provides evidence that calcium ions regulate heterocyst differentiation, Proc. Natl. Acad. Sci. U. S. A. 102, 16 (2005) 5744–5748.

[40] C. Steegborn, Structure, mechanism, and regulation of soluble adenylyl cyclases - similarities and differences to transmembrane adenylyl cyclases, Biochim. Biophys. Acta 1842, 12 Pt B (2014) 2535–2547. 10.1016/j.bbadis.2014.08.012

[41] N.M. Koropatkin, D.W. Koppenaal, H.B. Pakrasi, T.J. Smith, The structure of a cyanobacterial bicarbonate transport protein, CmpA, J. Biol. Chem. 282, 4 (2007) 2606–2614. 10.1074/jbc.M610222200

[42] I. Torrecilla, F. Leganes, I. Bonilla, F. Fernandez-Pinas, Use of recombinant aequorin to study calcium homeostasis and monitor calcium transients in response to heat and cold shock in cyanobacteria, Plant Physiol. 123, 1 (2000) 161–176. 10.1104/pp.123.1.161

[43] I. Torrecilla, F. Leganes, I. Bonilla, F. Fernandez-Pinas, Light-to-dark transitions trigger a transient increase in intracellular Ca2+ modulated by the redox state of the photosynthetic electron transport chain in the cyanobacterium Anabaena sp PCC7120, Plant Cell Environ. 27, 7 (2004) 810–819. DOI 10.1111/j.1365-3040.2004.01187.x

[44] I. Torrecilla, F. Leganes, I. Bonilla, F. Fernandez-Pinas, A calcium signal is involved in heterocyst differentiation in the cyanobacterium *Anabaena* sp. PCC7120, Microbiology 150, Pt 11 (2004) 3731–3739. 10.1099/mic.0.27403-0

