## Supplemental Figures for "Structural basis for the allosteric regulation of the SbtA bicarbonate transporter by the P_II_-like protein, SbtB, from *Cyanobium* sp. PCC7001"

**AMP (n=3)**

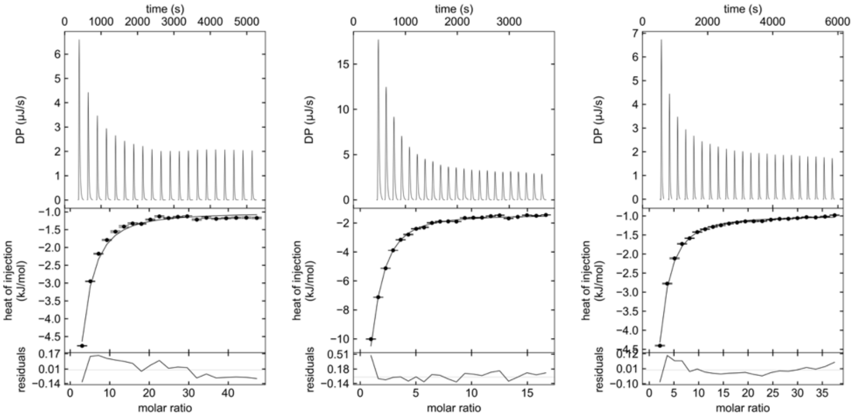


**Supp. Figure 1. ITC isotherms of titration of AMP into SbtB7001. (L-R) 1.** Cell = 102 μM SbtB7001 (monomer concentration) ; syringe =  20 mM AMP **2.** Cell = 400 μM SbtB7001 (monomer concentration); syringe =  23 mM AMP **3.** Cell = 143 μM SbtB7001 (monomer concentration) ; syringe =  20 mM AMP.

**AMP + 500 μM $CaCl_2$ (n=2)**

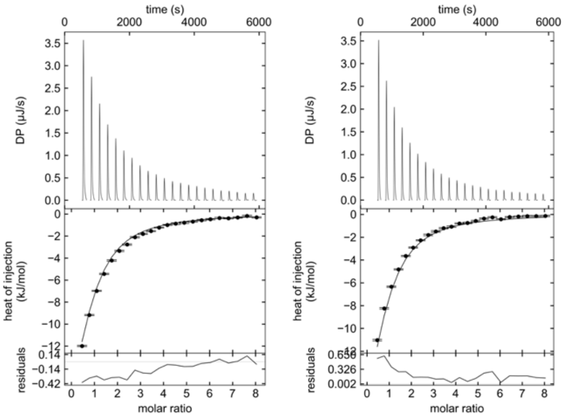


**Supp. Figure 2. ITC isotherms of titration of AMP into SbtB7001 in presence of $CaCl_2$. (L-R) 1.** Cell = 160 μM SbtB (monomer concentration) + 500 μM $CaCl_2$; syringe =  4 mM ATP + 500 μM $CaCl_2$ **2.** Cell = 160 μM SbtB (monomer concentration) + 500 μM $CaCl_2$; syringe =  4 mM ATP + 500 μM $CaCl_2$

**ADP (n=4)**

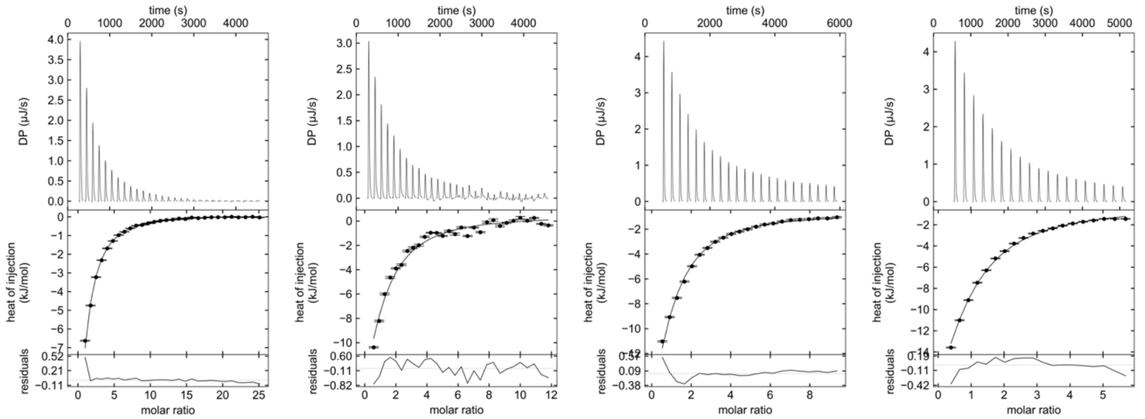


**Supp. Figure 3. ITC isotherms of titration of ADP into SbtB7001. (L-R) 1.** Cell = 102 μM SbtB7001 (monomer concentration); syringe = 10 mM ADP **2.** Cell = 110 μM SbtB7001 (monomer concentration); syringe = 5 mM ADP **3.** Cell = 143 μM SbtB7001 (monomer concentration); syringe = 5 mM ADP **4.** Cell = 160 μM SbtB7001 (monomer concentration); syringe = 4 mM ADP

**ADP + 500 μM $CaCl_2$ (n=2)**

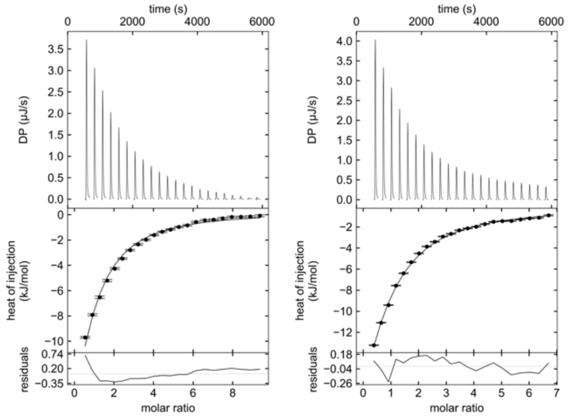


**Supp. Figure 4. ITC isotherms of titration of ADP into SbtB7001 in presence of $CaCl_2$. (L-R) 1.** Cell = 143 μM SbtB7001 (monomer concentration) + 500 μM $CaCl_2$; syringe = 5 mM ADP + 500 μM $CaCl_2$ **2.** Cell = 160 μM SbtB7001 (monomer concentration) + 500 μM $CaCl_2$; syringe = 4 mM ADP + 500 μM $CaCl_2$

**cAMP (n=4)**

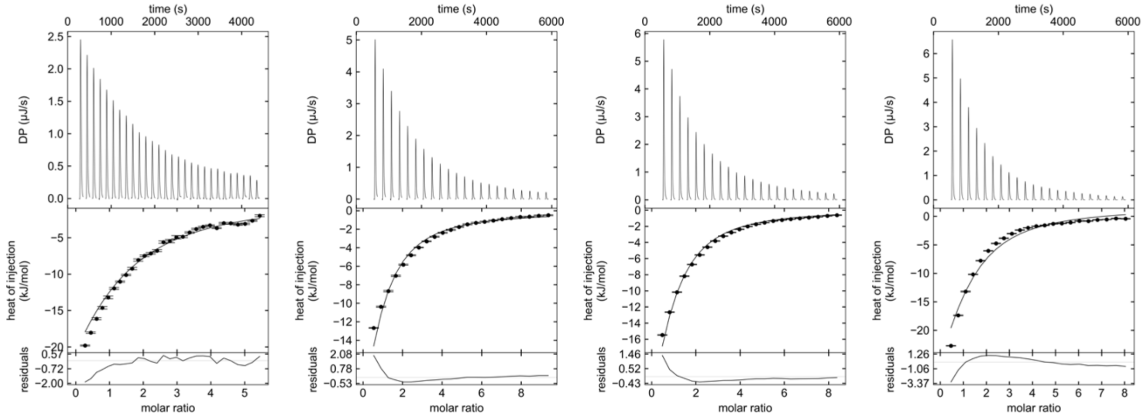


**Supp. Figure 5. ITC isotherms of titration of cAMP into SbtB7001. (L-R) 1.** Cell = 110 μM SbtB7001 (monomer concentration); syringe = 2 mM cAMP **2.** Cell = 143 μM SbtB7001 (monomer concentration); syringe = 5 mM cAMP **3.** Cell = 160 μM SbtB7001 (monomer concentration); syringe = 5 mM cAMP **4.** Cell = 160 μM SbtB7001 (monomer concentration); syringe = 4 mM cAMP

**cAMP + 500 μM $CaCl_2$ (n=2)**

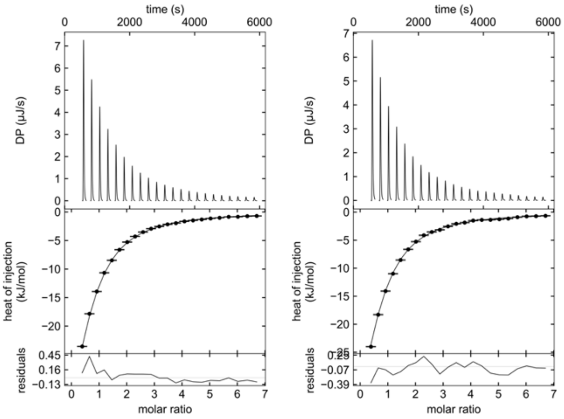


**Supp. Figure 6. ITC isotherms of titration of cAMP into SbtB7001 in presence of $CaCl_2$. (L-R) 1.** Cell = 160 μM SbtB7001 (monomer concentration) + 500 μM $CaCl_2$; syringe = 4 mM cAMP + 500 μM $CaCl_2$. **2.** Cell = 160 μM SbtB7001 (monomer concentration) + 500 μM $CaCl_2$; syringe = 4 mM cAMP + 500 μM $CaCl_2$.

**ATP (n=5)**

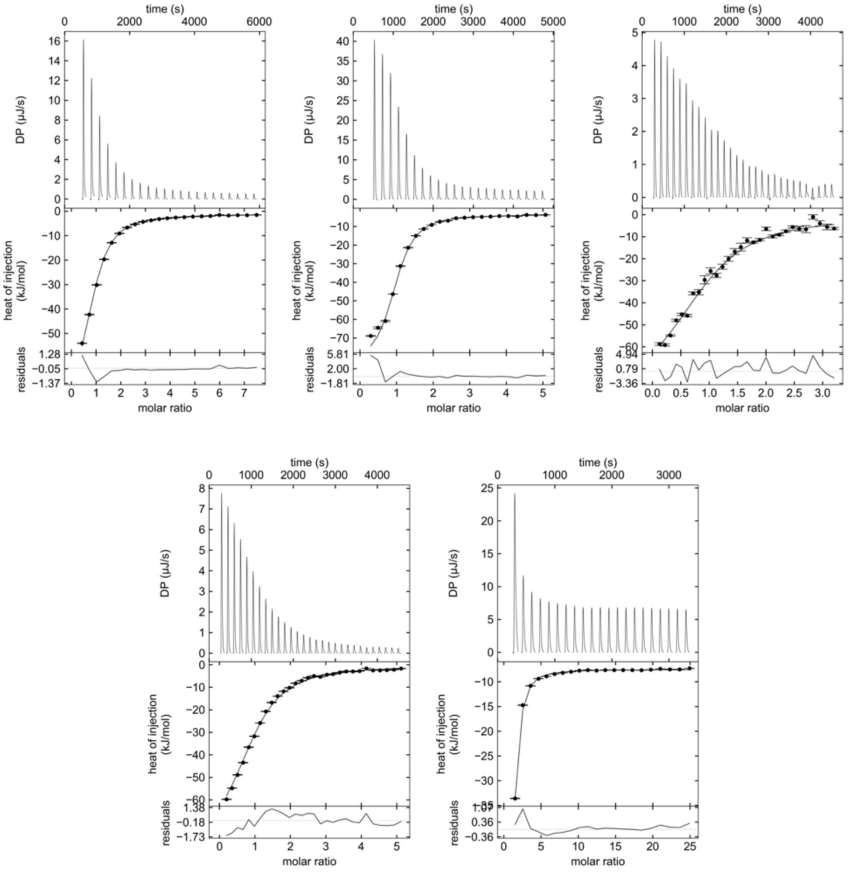


**Supp. Figure 7. ITC isotherms of titration of ATP into SbtB7001. (L-R) 1.** Cell = 143 μM SbtB7001 (monomer concentration); syringe = 4 mM ATP **2.** Cell = 400 μM SbtB7001 (monomer concentration); syringe = 7 mM ATP **3.** Cell = 96 μM SbtB7001 (monomer concentration); syringe = 1.2 mM ATP **4.** Cell = 100 μM SbtB7001 (monomer concentration); syringe = 2 mM ATP **5.** Cell = 102 μM SbtB7001 (monomer concentration); syringe = 10 mM ATP

**ATP + 10 mM $MgCl_2$ (n=2)**

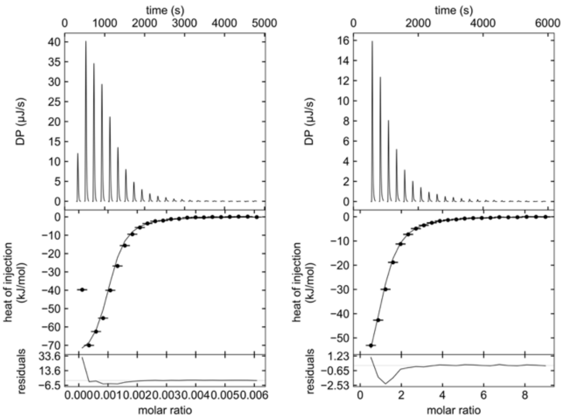


**Supp. Figure 8. ITC isotherms of titration of ATP into SbtB7001 in the presence of $MgCl_2$. (L-R) 1.** Cell = 400 μM SbtB7001 (monomer concentration) + 10 mM $MgCl_2$; syringe = 4 mM ATP + 10 mM $MgCl_2$ **2.** Cell = 143 μM SbtB7001 (monomer concentration) + 10 mM $MgCl_2$; syringe = 4 mM ATP + 10 mM $MgCl_2$

**ATP + EDTA**

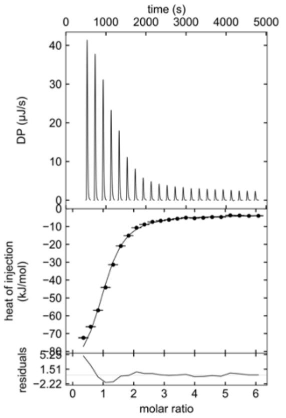


**Supp. Figure 9. ITC isotherm of titration of ATP into SbtB7001 in the presence of EDTA.** Cell = 400 μM SbtB7001 + 0.5 mM EDTA; syringe = 7 mM ATP + 0.5 mM EDTA

**ATP + 7 mM $CaCl_2$ (n=3)**

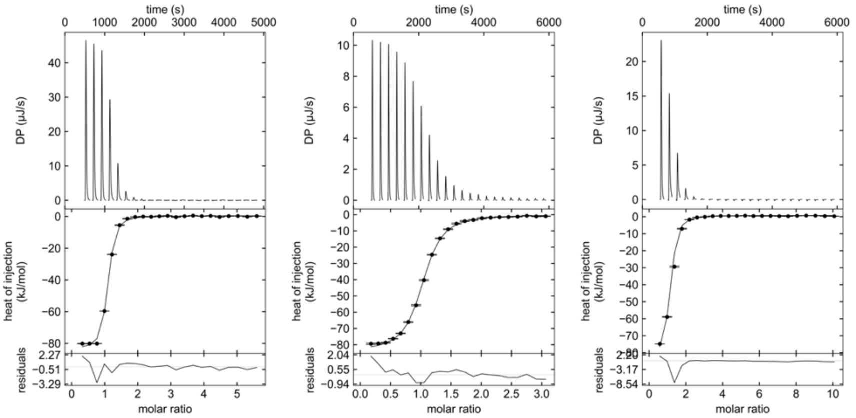


**Supp. Figure 10. ITC isotherms of titration of ATP into SbtB7001 in the presence of 7 mM $CaCl_2$. (L-R) 1.** Cell = 400 µM SbtB7001 (monomer concentration) + 7 mM $CaCl_2$; syringe = 7 mM ATP + 7 mM $CaCl_2$ **2.** Cell = 143 µM SbtB7001 (monomer concentration) + 7 mM $CaCl_2$; syringe = 1.5 mM ATP + 7 mM $CaCl_2$ **3.** Cell = 143 µM SbtB7001 (monomer concentration) + 7 mM $CaCl_2$; syringe = 4 mM ATP + 7 mM $CaCl_2$

**ATP + 500 µM $CaCl_2$ (n=2)**

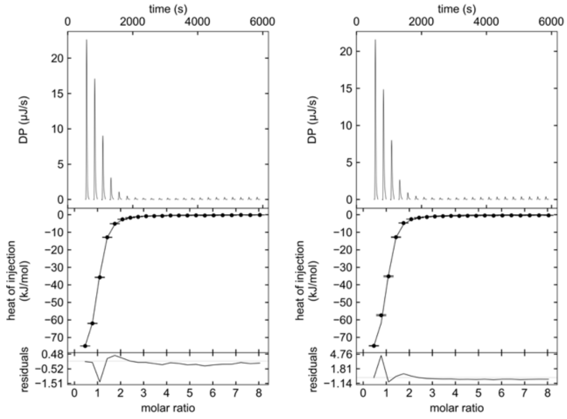


**Supp. Figure 11. ITC isotherms of titration of ATP into SbtB7001 in the presence of 500 µM $CaCl_2$. (L-R) 1.** Cell = 160 µM SbtB7001 (monomer concentration) + 500 µM $CaCl_2$; syringe = 4 mM ATP + 500 µM $CaCl_2$ **2.** Cell = 160 µM SbtB7001 (monomer concentration) + 500 µM $CaCl_2$; syringe = 4 mM ATP + 500 µM $CaCl_2$

**$CaCl_2$ (n=1)**

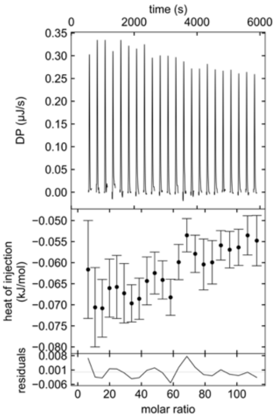


**Supp. Figure 12. ITC isotherm of titration of $CaCl_2$ into SbtB7001.** Cell = 143 μM SbtB7001 (monomer concentration); syringe = 50 mM $CaCl_2$

**$CaCl_2$ + 400 μM ATP (n=3)**

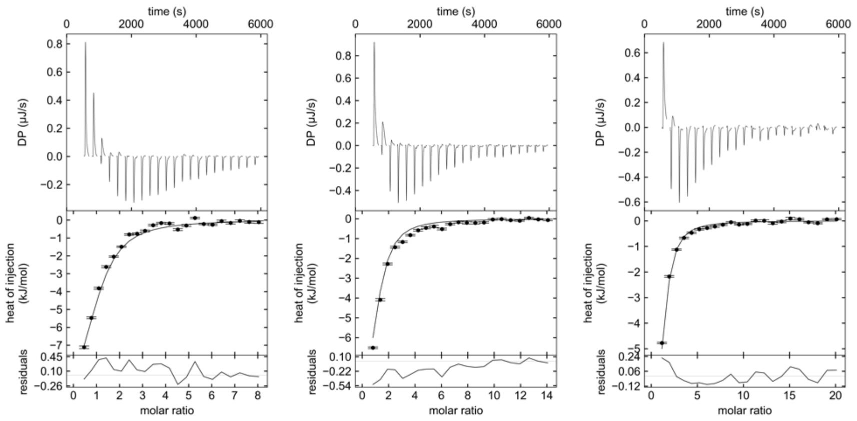


**Supp. Figure 13. ITC isotherms of titration of $CaCl_2$ into SbtB7001 in the presence of ATP. (L-R) 1.** Cell = 160 μM SbtB7001 (monomer concentration) + 400 μM ATP; syringe = 4 mM $CaCl_2$ + 400 μM ATP **2.** Cell = 160 μM SbtB7001 (monomer concentration) + 400 μM ATP; syringe = 7 mM $CaCl_2$ + 400 μM ATP **3.** Cell = 160 μM SbtB7001 (monomer concentration) + 400 μM ATP; syringe = 10 mM $CaCl_2$ + 400 μM ATP

**$NaHCO_3$ (n=2)**

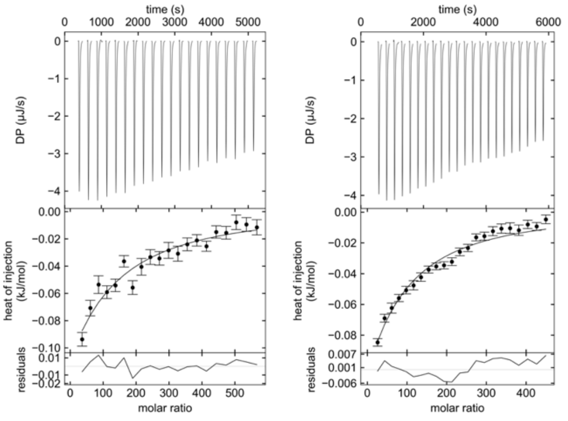


**Supp. Figure 14. ITC isotherms of titration of $NaHCO_3$ into SbtB7001. (L-R) 1.** Cell = 102 μM SbtB7001 (monomer concentration); syringe = 20 mM $NaHCO_3$ **2.** Cell = 143 μM SbtB7001 (monomer concentration); syringe = 200 mM $NaHCO_3$

**$NaHCO_3$ into LacI (n=2)**

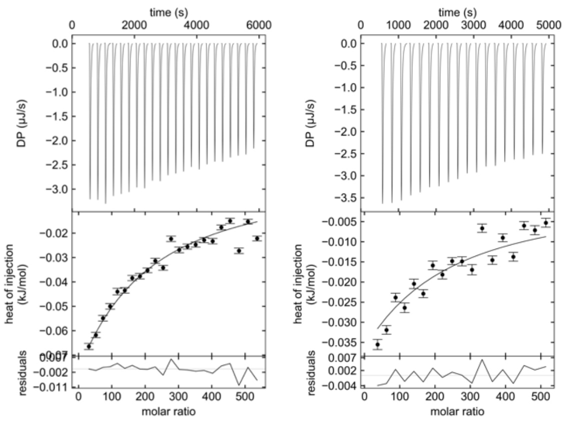


**Supp. Figure 15. ITC isotherms of titration of $NaHCO_3$ into *E. coli* lacI. (L-R) 1.** Cell = 100 μM *E. coli* lacI; syringe = 200 mM $NaHCO_3$ **2.** Cell = 100 μM *E. coli* lacI; syringe = 200 mM $NaHCO_3$
